# Maize associated bacterial and fungal microbiomes show contrasting conformation patterns that are resilient to water availability

**DOI:** 10.1101/2024.05.14.594156

**Authors:** Sandra Díaz-González, Sara González-Bodí, Carlos González-Sanz, Patricia Marín, Frédéric Brunner, Soledad Sacristán

## Abstract

Plant-associated microorganisms can help crops to alleviate water stress and increase the resilience of agricultural ecosystems to climate change. However, we still lack knowledge on the dynamics of bacterial and fungal microbial kingdoms within the soil and plant microbiomes and the response of these communities to different conditions such us, for example, water restrictions. This information is essential for the development of microbiome-based solutions to improve crop resilience to stressors associated to climate change. In this work, we explored: i) the conformation of the bacterial and fungal assemblages of different soil and plant compartments (bulk soil, rhizosphere, roots, leaves and grains) along the crop cycle of maize in an open field trial; and ii) the effect of water restriction on the maize microbiome comparing optimal irrigation with a 30% reduction of water supply. Our results show that microbial communities are highly structured along soil and plant compartments, with contrasting patterns for bacteria and fungi that were intensified towards the end of the plant cycle. Root showed the most differentiated bacterial assemblage while fungi conformed a very distinct community in the leaf, suggesting a relevant contribution of aerial fungal propagules to the microbiome of this plant organ. Despite the reductions in plant growth and yield, the microbiome of limited-watered plants did not show severe alterations. Still, significant impacts were observed within compartments, being fungi more responsive to limited watering than bacteria. Network analysis suggest that bacteria and fungi may play different roles in the shifts observed under water stress.

## 1. Introduction

Plant microbiomes are composed of a huge variety of microorganisms, including bacteria, fungi, oomycetes, archaea and viruses, which directly participate in relevant functions of the plant (Vandenkoornhuyse et al., 2015; Rodriguez et al., 2019), such as seed germination, plant development, stress tolerance, nutrient uptake, crop productivity or synthesis of relevant bioactive compounds (Bulgarelli et al., 2013) The conformation of the plant microbiome relies on a dynamic and selective recruitment of microorganisms by the host during its development, which is also influenced by external abiotic and biotic constraints (Hassani et al., 2019; Mesny et al., 2023). Microbe-microbe interactions may also play a critical role in plant fitness and its response and adaptation to the external environment.

Despite research on plant microbiota has grown exponentially in the last decade, most of the published works have focused on one particular microbial kingdom, mostly bacteria, and one or few host compartments (Hacquard, 2016). This fact has provided an extensive understanding of the functioning of the bacterial microbiome of specific compartments, such as the rhizosphere or the roots. However, we still lack a holistic view of the construction of the plant microbiome, its evolution with plant growth and its response to environmental stressors (Nilsson et al., 2019a; Mesny et al., 2023).

One important environmental stressor in agroecosystems is water scarcity. Over the next decades, variation and uncertainty in precipitation patterns among years and seasons are expected to increase due to climate change (IPCC, 2018, 2019). Many crops are highly sensitive to water stress, either by restrictions in irrigation or reduced precipitation (Daryanto et al., 2016; Song et al., 2019). Irrigation demand, which already holds a great part of global water usage, is expected to increase in the next decades (Wada et al., 2013). Furthermore, the quality and quantity of the water resources are decreasing, being water limitation one of the main concerns (Reilly et al., 2003; Sabale et al., 2022). Therefore, there is a need to understand the mechanisms and factors through which water stress affects crops and their surrounding ecosystem in order to bring solutions that mitigate the impact of water scarcity and preserve crop productivity and food security.

Limitation in water availability can have a substantial impact on plant associated microbial communities (Santos-Medellín et al., 2017; Cavicchioli et al., 2019; Williams and Vries, 2020; Bei et al., 2023). On the one hand, drought is very likely the abiotic stress with the largest direct repercussions on soil microbiota, the main reservoir of plant-associated microbes (De Vries et al., 2020). Evolution has provided these microorganisms with diverse strategies to overcome water stress, such as osmoregulation, sporulation, dormancy, extracellular enzyme synthesis and osmolyte production (Barnard et al., 2013). However, strong osmotic stress, soil heterogeneity and limited nutrient mobility produced by water scarcity result in a direct reduction of microbial soil biomass which affects the capacity of soil to support plant growth and development (Jansson and Hofmockel, 2020). On the other hand, plants react to water stress by triggering a cascade of metabolic and physiological processes which also influence the assembly of the plant microbiome (Xu et al., 2010; Chen et al., 2022).

In this work we have studied the microbiome of maize plants in an open field trial where two different irrigation regimes were applied: and optimal irrigation program and a 30% of water restriction from the optimal irrigation regime. Our initial hypothesis is that plant microbiome conformation along soil-plant compartmental niches follows a robust pattern, but it can be modulated by environmental conditions, such as water availability. Thus, with this study we aim to answer the following questions: i) how are maize-associated bacterial and fungal microbiomes conformed along the plant cycle?; ii) to what extent does water availability affect plant-associated microbiome conformation? and iii) do plant associated bacterial and fungal assemblages respond differently to water availability? For that, we have first carried out a detailed description of the microbiome conformation of maize associated bacterial and fungal communities in five different soil and plant compartments (bulk soil, rhizosphere, roots, leaves, and grains) along three time points of plant cycle (one, two and four months after sowing) in our open field trial and, second, we have analysed the effect of water limitation in the bacterial and fungal assemblages, as well as in the network interactions within and between these two microbial kingdoms. Our results confirm the strong compartmentalization of plant associated microbiome, showing a dynamic compartment-driven recruitment of microorganisms that is intensified at the end of the plant cycle, with differential patterns for bacteria and fungi. The impact of water availability was rather observed within soil and plant compartments and was stronger in fungal assemblages, where we can define hallmark fungal ASVs for each compartment and irrigation regime. Bacteria and fungi seemed to have different roles in the topological properties shifts of microbial networks under water stress conditions.

## 2. Material and methods

### 2.1. Site description, plant material and trial design

The study was conducted from May to October 2018 in the field of an experimental station located at Pilar de la Horadada, Alicante (37°51’46.0”N-0°48’30.6”W, 35 m above mean sea level) in the south-eastern part of the Iberian Peninsula. According to Köppen – Geiger classification, the climate in this area is hot steppe (BSh) (AEMET 2011) with average annual temperature of 17.6°C (ranging from 10.8°C in January to 25.5°C in August). Mean annual precipitation is 313 mm, with June, July, and August being especially dry months (average of 7, 2, and 7 mm respectively; AEMET, 2010). The mean annual humidity is 71% (AEMET, 2010). Thus, in order to meet maize water requirements in this area, irrigation is needed.

The maize commercial variety LG 34.90 (Limagrain Ibérica SA) was used in this work. Soil fertilization was adjusted based on maize nutritional needs and soil physical-chemical analyses as previously described in (Díaz-González et al., 2020). The experimental area was divided into two blocks of 4 plots each: one block received 100% of the water requirements (optimal watering, OW), while the other had a water restriction of 30%, so received 70% of optimal water requirements (limited watering, LW) (Supplementary Figure 1). Water restriction started two weeks after sowing to ensure optimal seed germination. Each plot consisted in three rows of 9 m and a total area of 20.25 m^2^. The total area of the trial was 324 m^2^.

### 2.2. Plant growth and yield measurements

Stem height and Chlorophyll Content Index (CCI) were recorded from 20 plants per plot (80 plants per irrigation regime) from the middle row of each plot at reproductive stage R4 (dough stage, milky inner fluid of kernels thickens to a pasty consistency)(Nleya et al., 2019). Stem height was determined from the ground to the beginning of the male inflorescence.

To calculate mean cob weight per plant, mature cobs from 44 plants from the same plot were collected and weighted together. Cobs were dekernelled and grains were weighted together to determine mean plant yield for each plot. Grain humidity was assessed in order to normalize weights to 15.5% of water content. Yield in t/ha was calculated according to trial plant density (75.000 plants/ha).

### 2.3. Statistical analysis of vegetative and productive parameters

Statistical analyses were performed in RStudio software (version 4.2.2). In order to check normality and homogeneity of variances assumptions, both Shapiro – Wilk Normality Test and Bartlett test of Homogeneity of Variance were performed using ‘shapiro.test’ and ‘bartlett.test’ functions, respectively (R package stats). Two-sample comparisons for data fitting to a normal distribution were analysed by Welch Two-sample t-test with ‘t.test’ (*P* < 0.05) function (R package stats). In two-sample comparisons not meeting normality assumption a Two-Sample Wilcoxon Test (*P* < 0.05) was conducted by the implementation of ‘wilcox.test” function (R package stats).

### 2.4. Sample collection, processing, and DNA extraction for maize microbiome analysis

In order to have a representation of the whole trial area, we defined the plot as our sample unit. One maize plant per each OW or LW plots (4 plants per irrigation regime, 8 plants in total, see Supplementary Figure S1) were randomly selected and collected at three time points: i) T01: one month after sowing (maize early developmental stage V4-5); ii) T02: two months after sowing (mid maize developmental stage V8-9); iii) T03: near to the end of the crop cycle (reproductive stage R4; approximately 4 months after sowing) (Nleya et al., 2019). From each plant, bulk soil, rhizosphere, root, leaf, and grain compartments were collected (n = 24 per compartment, except grain). Grains were only collected in T03 (n = 8). In total, 104 samples were processed as specified in Supplementary Methods.

Total genomic DNA was extracted from bulk soil, rhizosphere, and grain samples with FastDNA^TM^ Spin Kit for Soil (MP Biomedicals) following manufacturer’s instructions. DNA extraction from root and leaf tissues was conducted with DNeasy Plant Mini Kit (Qiagen), following manufacturer’s instructions with a minor adaptation in elution volumes (30 µL instead of 100 µL). Centrifugation steps were carried out at 4°C. DNA quality was verified by agarose electrophoresis (1.5% agarose, Agarose D1 Medium EEO, Conda Pronadisa). DNA quantification was carried out with Quant-iT^TM^ PicroGreen^TM^ dsDNA Assay Kit (Invitrogen) in Varioskan^TM^ LUX Multimode microplate reader (Thermo Scientific).

### 2.5. Amplicon generation and sequencing

Fifty µL of each DNA extract were sent to Sequentia Biotech S.L. for DNA normalization, amplicon generation, library preparation, and sequencing. Bacterial and fungal amplicons were obtained by PCR from DNA extracts of bulk soil, rhizosphere, root, leaf, and grain samples by using primer pair 515f/806r (Caporaso et al., 2011) for bacterial 16S rRNA gene and ITS1-F (Gardes and Bruns, 1993)/ITS2 (White et al., 1990) for fungal ITS gene. Amplicon simultaneous sequencing was conducted on Illumina’s HiSeq2500 platform, generating 2x250bp reads.

### 2.6. Read data processing and taxonomic assignment

The Divisive Amplicon Denoising Algorithm 2 (DADA2) (version 1.22.0) was employed for the taxonomic identification of bacterial and fungal amplicons. The default parameters were used, with certain exceptions as specified (Callahan et al., 2016). Initially, all raw reads were processed for adapter trimming using the cutadapt tool (version 4.6) (Martin, 2011). For the 16S rRNA gene sequencing data, standard filtering parameters were applied. These parameters allowed no uncalled bases, a maximum of 2 expected errors, and truncated reads at a quality score of 2 or less. In the case of the fungal ITS dataset, the same filtering approach was applied, with an additional parameter specifying a minimum read length of 50 bp. In both instances, learned errors were utilized to infer predicted error presence across all reads as a denoising measure.

Amplicon Sequence Variants (ASVs), where each ASV differs from the others by at least one nucleotide, were identified in each sample after sequence de-replication. Chimeric ASVs were determined with the consensus method. Subsequently, the ASVs were mapped to the SILVA v138 database (Yilmaz et al., 2014) with a minimum bootstrap confidence level of 80 for bacterial assignment. The taxonomic assignment for ITS was performed against the UNITE v18.11.2018 database using the same bootstrap confidence level as above (Nilsson et al., 2019b).

After DADA2 processing, the datasets retained the following number of reads: 41,815,842 (65.67 % average among samples) for the 16S rRNA gene dataset, and 102,449,632 (77.49%) for the ITS dataset (Supplementary Table S1.1). The recovery of reads through both workflows was tracked for each step in each sample (Supplementary Table S1.2 and S1.3 respectively).

### 2.7. Bioinformatic analysis

Downstream analyses were performed using RStudio software (version 4.2.2) and R (version 4.3.2). The resulting abundance table and taxonomic classification generated by DADA2 were imported into the Phyloseq program (version 1.20.0) (McMurdie and Holmes, 2013). Plant reads were discarded. We filtered out sequence chimeras or artifacts by removing ASVs without class or order taxonomic assignment. ASVs that were low represented with fewer than three reads and prevalent in less than 2% of the total samples were also discarded. Then, taxa counts were normalized using median sequencing depth. We obtained 7,579,959 (16S rRNA gene) and 64,050,053 (ITS) high-quality reads that were assigned to 11,246 bacterial ASVs and 2,080 fungal ASVs (Supplementary Tables S1 and S2). Rarefaction curves showed that sequencing was satisfactory to accurately uncover sample diversity (Supplementary Figure S2).

#### Diversity analysis

Alpha-diversity (diversity within samples) analysis was carried out based on both Chao1 richness estimator and Shannon index at ASV level with the function ‘estimate_richness’ (R package phyloseq). Kruskal-Wallis Rank Sum Test (*P* < 0.05) was undertaken for Chao1 and Shannon index variations among compartments and time points using ‘kruskal.test’ function (R package stats). Two-Sample Wilcoxon Test (*P* < 0.05) was conducted by the implementation of ‘wilcox.test’ function (R package stats) for the comparison between irrigation regimes. Beta-diversity analysis was carried out by calculating Bray-Curtis dissimilarity at the ASV level by using functions ‘distance’ and ‘ordinate’ (R package phyloseq). Permutational multivariate analyses of variance (PERMANOVA) were performed using the ‘adonis2’ function implemented in the vegan package (version 2.6.4). Plots were generated with ‘ggplot’ function (R package ggplot2).

#### Taxonomic composition

Taxa composition was calculated at the genus level. ASVs were merged to genus level to determine relative abundance. Plots were generated with ‘ggplot’ function (R package ggplot2).

#### Venn diagrams and heatmaps construction

R function ‘ps_venn’ (R_package MicEco) was used to extract the ASVs that overlapped among compartments, while ‘venn.diagram‘ (R package VennDiagram) function was used for diagram plotting. Heatmaps were constructed based on the list of ASVs that were shared by at least two compartments. Function ‘Heatmap’ from R package ComplexHeatmap was used to visualize the graph. Cluster evenness was calculated with function ‘evenness’ from R package tabula. Cluster taxonomic composition was calculated as mentioned above and pie charts were created with GraphPad Prism (version 8.0.1).

#### Network analyses

SparCC algorithm (Friedman and Alm, 2012) implemented with FastSpar (Watts et al., 2019) was used to infer co-occurrence patterns at the ASV level. Only ASVs present in at least 25% of the samples with total counts >10 were used for network construction and analysis. The networks were visualized and analysed using Cytoscape 3.10.1 (Shannon et al., 2003). In the networks, nodes represent interconnected ASVs and edges the connections (positive or negative co-occurrence) among nodes. Analysis was conducted on nodes and edges with high and significant Spearman’s correlation scores (*r* > 0.6 or *r* < − 0.6; *P* < 0.05). The degree is the number of edges of a specific node. Betweenness centrality is a measure of the extent of control exerted by a given node over the interactions among other nodes in the network (Yoon et al., 2006). Network hubs were defined as nodes with values of degree and betweenness centrality higher or equal to percentile 90^th^ (≥ 𝑃_90_) within each network. Pie charts showing the proportion of inter- and intra-kingdom positive and negatives edges were created with GraphPad Prism (version 8.0.1). Correlations between number of ASVs, nodes, edges and number of hubs were calculated with R function ‘rcorr’ from package Hmisc.

## 3. Results

### 3.1. Limited irrigation had a negative impact on maize growth and yield

We measured vegetative parameters at phenological stage R4. OW plants were significantly taller (*P* < 0.001) and showed significant higher shoot biomass (*P* < 0.001) than LW plants (Table 1). LW plants also showed a significant reduction in CCI (*P* < 0.001). Reproductive and yield parameters were measured at the end of the plant cycle. LW plants showed a significant decrease in mean cob weight per plant (*P* = 0.002) that affected yield, resulting in a significant reduction of almost 5 t/ha (39.2% reduction; *P* < 0.001) compared with OW plants. The yield obtained in OW plants was comparable to the expected according to the data provided by the seed breeder, similar to the average yield of irrigated maize in Spain (12.3 t/ha) and 1.5 tons higher than the average in the province of Alicante (11 t/ha) in 2018 (Ministerio de Agricultura Pesca y Alimentación, 2019). Therefore, OW plants attained the expected optimal growth and the applied water restriction had a negative impact in several vegetative and productive parameters of maize plants.

**Table 1.** Growth and yield of maize plants subjected to optimal (OW) or limited (LW) watering.

| Irrigation | CCI | Stem height (m) | Shoot fresh weight (g) | Cob weight per plant (g) | Yield (t/ha) |
| --- | --- | --- | --- | --- | --- |
| OW | 45.0 ± 11.2 a | 2.5 ± 0.24 a | 599.9 ± 138.0 a | 196.0 ± 21.5 a | 12.5 ± 1.9 a |
| LW | 36.4 ± 8.4 b | 2.1 ± 0.21 b | 452.2 ± 116.8 b | 151.0 ± 24.8 b | 7.6 ± 1.9 b |

Data from stem height (n=160) and CCI (n=160) were analysed by non-parametric Wilcoxon-Mann-Whitney test. Data for shoot fresh weight (n=32), cob weight per plant (n=8) and yield (n=8) were analysed by Student's T test. Different letters within each column indicate significant differences among treatments ( $P < 0.05$ ). Data show means ± standard deviations.

### 3.2. Soil and plant compartments are the main factors affecting microbiome richness and alpha-diversity with minor effect of irrigation regime

Alpha-diversity (diversity within samples) was measured by both Chao1 index richness estimator and Shannon index, which takes into account both species richness and evenness (Figure 1, Supplementary Table S3). Bacterial Chao1 and Shannon indexes were much higher than fungal (in the order of ten and two times, respectively) especially for soil compartments (bulk soil and rhizosphere). Indeed, richness was always much higher in soil than in plant compartments (Figure 1A). This was especially remarkable in the bacterial microbiome, where Chao1 index showed a severe reduction along the plant, being up to 38 times lower in the leaf than in the bulk soil. Thus, differences in richness between bacterial and fungal communities decreased strongly from soil to plant compartments and within the plant from roots to leaves, reaching very similar Chao1 indexes in the leaf in both (bacterial and fungal) microbiomes. Richness in plant compartments increased with time, resulting in a decrease in richness differences between soil and plant compartments for both bacterial and fungal microbiomes towards the end of the plant cycle.

**Figure 1:**
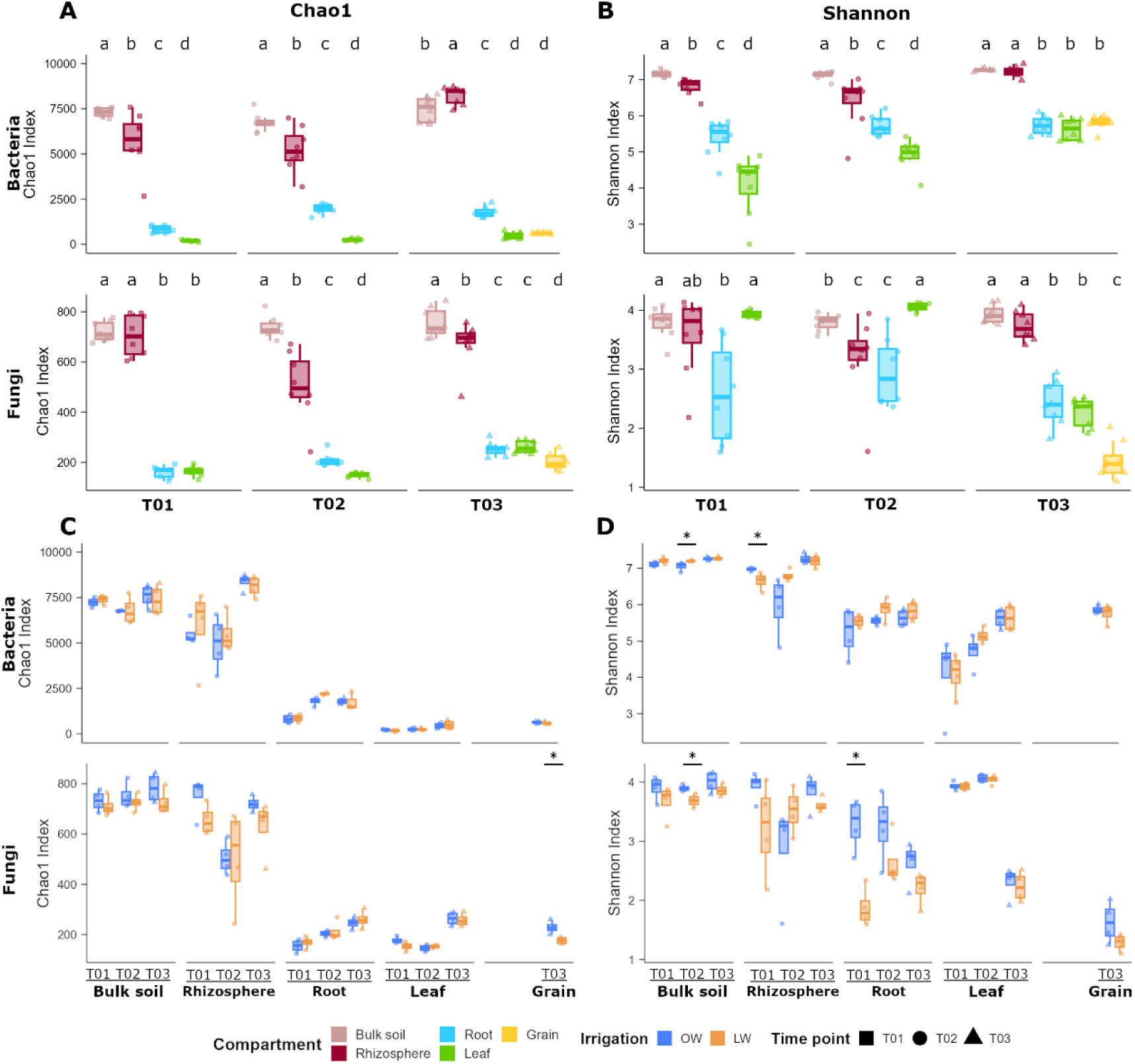
Alpha diversity dynamics of maize-associated microbiome at different compartments. A) Chao1 and B) Shannon indexes of bacterial and fungal communities in different compartments (bulk soil, rhizosphere, root, leaf and grain) during different time points (one -T01-, two -T02- and four -T03- months after sowing) of maize cycle. C) Chao1 and D) Shannon indexes of bacterial and fungal communities in different compartments during different time points (one -T01-, two -T02- and four -T03- months after sowing) and irrigation regimes (OW: optimal watering, LW: 30% watering reduction). Internal line in box-plot boxes indicates the median or second quartile (Q2), upper line of the boxes indicates the third quartile (Q3) and lower line the first quartile (Q1) of the data. Different letters and asterisks above the boxes indicate significant differences determined by non-parametric Wilcoxon-Mann-Whitney Test.

Shannon indexes were also higher in soil than in plant compartments, but differences between rhizosphere and soil and between leaf and root were gradually reduced with time, being similar at T03 (Figure 1B). In the fungal assemblage, however, this index was very similar among compartments at earlier time points and tended to differentiate at the end of the cycle (T03), when we observed a steady decrease in diversity from bottom to top-plant compartments (Figure 1B). Thus, the increase in fungal richness with time in plant compartments (Figure 1A) was not reflected in a higher Shannon index, that did not change in the root (*P* = 0.229) or even decreased significantly in the leaf (*P* < 0.001) (Figure 1B). This increase in the number of fungal taxa towards the end of the cycle joint to a decline in diversity was due to a preponderance of specific taxa in the leaves that negatively affected the evenness (Supplementary Figure S3). Seven out of the ten most abundant genera showed an abrupt decrease in relative abundance at T03. By example, *Exserohilum* shifted from the top 1 with 13.2% of abundance at T01 to 0.12% at T03 (*P* <0.001). In contrast, the genera *Alternaria* and *Cladosporium,* which together accounted for 84% of the relative abundance at T03, did not reach 20% at earlier stages.

When analysing diversity separately by irrigation regime, similar trends were observed, with mostly no significant differences (Figure 1C and D, Supplementary Table S3). However, LW tended to negatively affect richness and Shannon index of fungal assemblages. Thus, when aggregating all the datasets per water regime, the Shannon index of fungal assemblages was significantly lower in LW than OW microbiomes (*P* = 0.042, Supplementary Table S3). When looking at compartments and time points separately, the significant reductions linked to water restriction in fungal microbiomes were observed for Shannon index at bulk soil (T02) and root (T03), and Chao1 index for grain (T03).

### 3.3. Conformation of microbial assemblages along the plant is different for bacteria and fungi

The impact of the different factors on beta-diversity (or diversity among samples) was analysed by means of Bray-Curtis dissimilarity Principal Coordinate analysis (PCoA) and PERMANOVAs (Figure 2, Supplementary Figure S4 and Supplementary Table S4). These analyses showed that compartment was the main driver of microbiome differentiation (Bacteria: R^2^= 0.385, *P* < 0.001; Fungi: R^2^ = 0.448, *P* < 0.001), followed by time (Bacteria: R^2^= 0.066, *P* < 0.001; Fungi: R^2^= 0.075, *P* < 0.001). Moreover, the interaction between compartment and time was significant for both bacteria (R^2^= 0.133, *P* < 0.001) and fungi (R^2^= 0.139, *P* < 0.001), meaning that the time did not affect equally to microbiome composition in each compartment. Irrigation regime was not significant in PERMANOVA analysis on the aggregated dataset (Bacteria: R^2^= 0.015, *P* = 0.111; Fungi: R^2^= 0.018, *P* = 0.057). Nonetheless, the interaction between compartment and irrigation regime was significant for the fungal assemblage (R^2^= 0.036, *P* = 0.014).

**Figure 2:**
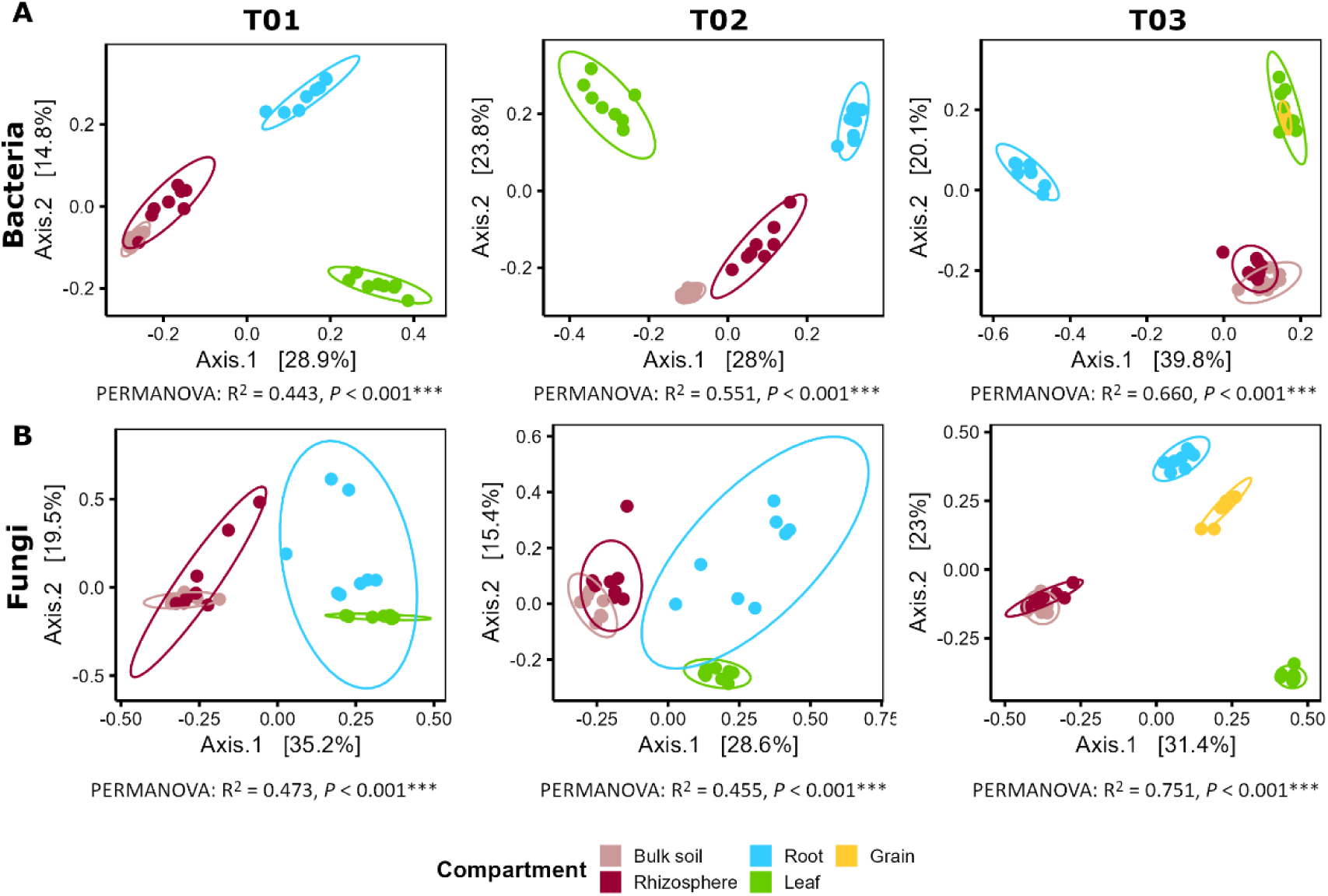
Beta-diversity patterns of maize-associated microbiome at different time points. Principal coordinate analysis (PCoA) based on Bray-Curtis dissimilarity depicting the distribution patterns of bacterial (A) and fungal (B) communities at different time points (one -T01-, two -T02- and four -T03- months after sowing) and unconstrained by compartment. The relative contribution (R^2^) of the factor “compartment” on community dissimilarity and its significance was tested with PERMANOVA. T01 and T02: n = 16; T03: n = 20.

The effect of time was significant in the conformation of both bacterial and fungal communities within each compartment, although with some differences. The proportion of variance explained by the time of sampling in the bacterial assemblage of belowground compartments, with 44.8% in the bulk soil, 35.6% in the rhizosphere and 43.9% in the root, was noticeably higher than in the fungal assemblage, with proportions of 14.4%, 22.5% and 21.7% in the bulk soil, rhizosphere and root, respectively (Supplementary Table S4). However, these proportions were inverted in the leaf, where the time accounted for 74.9% of the explained variance in the fungal assemblage, while only 21.3% was explained by this variable in the bacterial assemblage.

Figure 2 shows the microbiome differentiation at the different time points along soil and plant compartments. In the bacterial assemblage, we found a correspondence between physical distance and sample dissimilarity among compartments at earlier time points (T01 and T02). Samples from soil compartments (bulk soil and rhizosphere) grouped together and closer to root samples, while leaf samples were more dissimilar to the belowground samples and thus, more distantly located in the PCoA plots. However, towards the end of the cycle, at T03, samples from soil compartments got closer to the group formed by leaf and grain samples, becoming the root the most differentiated compartment along axis 1, which explains more than 40% of the variance (Figure 2A). In the case of fungi, we observed certain overlaps among compartments at earlier time points, but a very clear separation at T03, except for soil compartments that grouped together. In contrast to bacterial assemblage, the leaf was the most differentiated compartment in this case (Figure 2B). It is also noteworthy, that the composition of the microbiota in the grain was more similar to the leaf in the case of the bacterial assemblage (Figure 2A), and remarkably more similar to the root in the case of the fungal assemblage (Figure 2B).

One important factor for microbiome differentiation is the proportion of taxa common or exclusive to different samples. Venn diagrams show a core microbiome of 741 bacterial ASV and 241 fungal ASV present in all compartments, which account for 7% and 12% of full bacterial and fungal assemblages, respectively (Figure 3A and D, Supplementary Table S5). The rhizosphere was the compartment with the highest number of shared ASVs, with 9290 bacterial ASVs (83% of the total and 92% of rhizosphere ASVs) and 1711 fungal ASVs (82% of the total and 96% of the rhizosphere ASVs). Besides, soil and rhizosphere shared the largest group of ASVs, with 9041 bacterial ASV (97% of the 9290 shared ASVs of the rhizosphere) and 1613 fungal ASVs (94% of the 1711 shared ASVs of the rhizosphere). Compartment-exclusive represented only a small proportion of ASVs (Figure 3A and D, Supplementary Table S5). In soil compartments, the percentage of exclusive bacterial ASVs was higher than for fungi, but the opposite happened in plant compartments. Thus, in bacterial assemblages there was a considerable reduction in the proportion of exclusive ASVs from the soil compartments (bulk soil: 9.2% and rhizosphere: 8.4%) to the roots (1.9%), and from this to the aboveground compartments (leaf: 0.6% and grain: 0.5%). In fungi, the proportion of compartment-exclusive ASVs decreased from bulk soil (7.5%) to the roots (2.6%), growing in the leaves (7.5%) and being reduced again in the grain (4%) (Supplementary Figure S5 and Supplementary Table S5).

**Figure 3:**
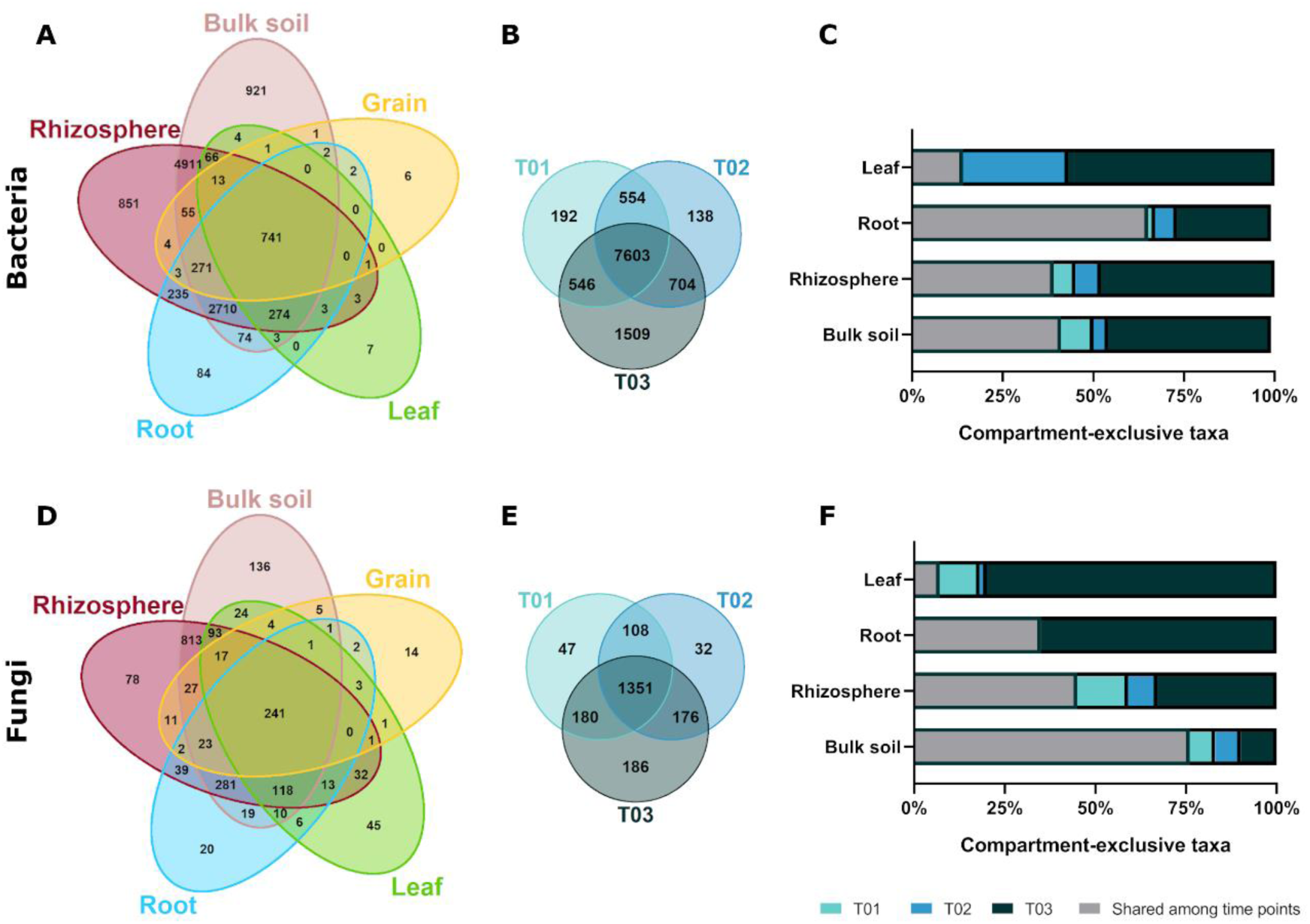
Configuration of maize-associated microbiome across plant compartments and time points. Venn diagrams showing exclusive and shared bacterial and fungal ASVs among (A and D) compartments and (B and E) time points (one -T01-, two -T02- and four -T03- months after sowing). Proportion of compartment exclusive bacterial (C) and fungal (F) ASVs exclusively present at T01, T02, T03 or present in more than two time points.

Most of the bacterial (7603 ASVs) and fungal (1351 ASVs) taxa, accounting for 67.6% and 65% respectively, were present during the whole plant cycle (Figure 3B and E; Supplementary Table S5). Time point T03, closer to the end of the plant cycle, was the one exhibiting the highest proportion of exclusive ASVs, with 1509 bacterial taxa (13.4%) and 186 fungal taxa (8.9%). The proportion of compartment-exclusive taxa increased towards the end of the cycle in all compartments (Figure 3C and F, Supplementary Table S6). However, the impact of time in the compartment-exclusive ASVs was different for bacteria and fungi. In bacteria, ASVs only present at T03 were predominant among bulk soil- and rhizosphere-exclusive ASVs (Figure 3C). In fungi, there was an increase from soil to leaves of ASVs only present at T03, being most of the compartment-exclusive taxa of roots and leaves only found at the latest time point (Figure 3F). These results suggest a specific and differential recruitment of bacterial and fungal taxa towards the end of the plant cycle, especially in the roots and the leaves.

### 3.4. Predominant fungal taxa are specific of plant compartment and irrigation regime

Given the strong effect of soil and plant compartments in the assemblage of maize-associated microbiome, we investigated if the irrigation regime could affect the assemblages within each compartment (Figure 4A, Supplementary Table S4). For bacterial microbiome, irrigation regime explained beta-diversity significantly only in the bulk soil and in the leaf, whereas for fungal microbiome it was significant in all the compartments except the leaf (Figure 4A, Supplementary Table S4). When looking to aggregated data, only a small proportion of the taxa were exclusively found in each irrigation regime (Figure 4B). In the bacterial assemblage, the percentages of exclusive ASV were very similar for each irrigation regime, with 738 exclusive ASVs (6.6%) for OW and 756 exclusive ASVs (6.7%) for LW. In the case of fungi, however, there was an enrichment of exclusive ASVs in OW plants, accounting for 9.7% (202 ASVs), in comparison to 5.6% (122 ASVs) in LW plants. When comparing data within each compartment, the proportion of ASVs exclusive for each water regime increases both in bacterial and fungal assemblages (Supplementary Table S5). The enrichment in exclusive ASVs within the compartments was more relevant in the fungal assemblage, being the root the compartment which exhibited a higher proportion of exclusive taxa, accounting for 20% (156 ASVs) in OW and 24% (187 ASVs) in LW (Supplementary Table S5). These results indicate that, although irrigation has a limited impact on the global recruitment of microbes by the plant, relevant changes can be accounted for within compartments, mainly in the fungal community.

**Figure 4:**
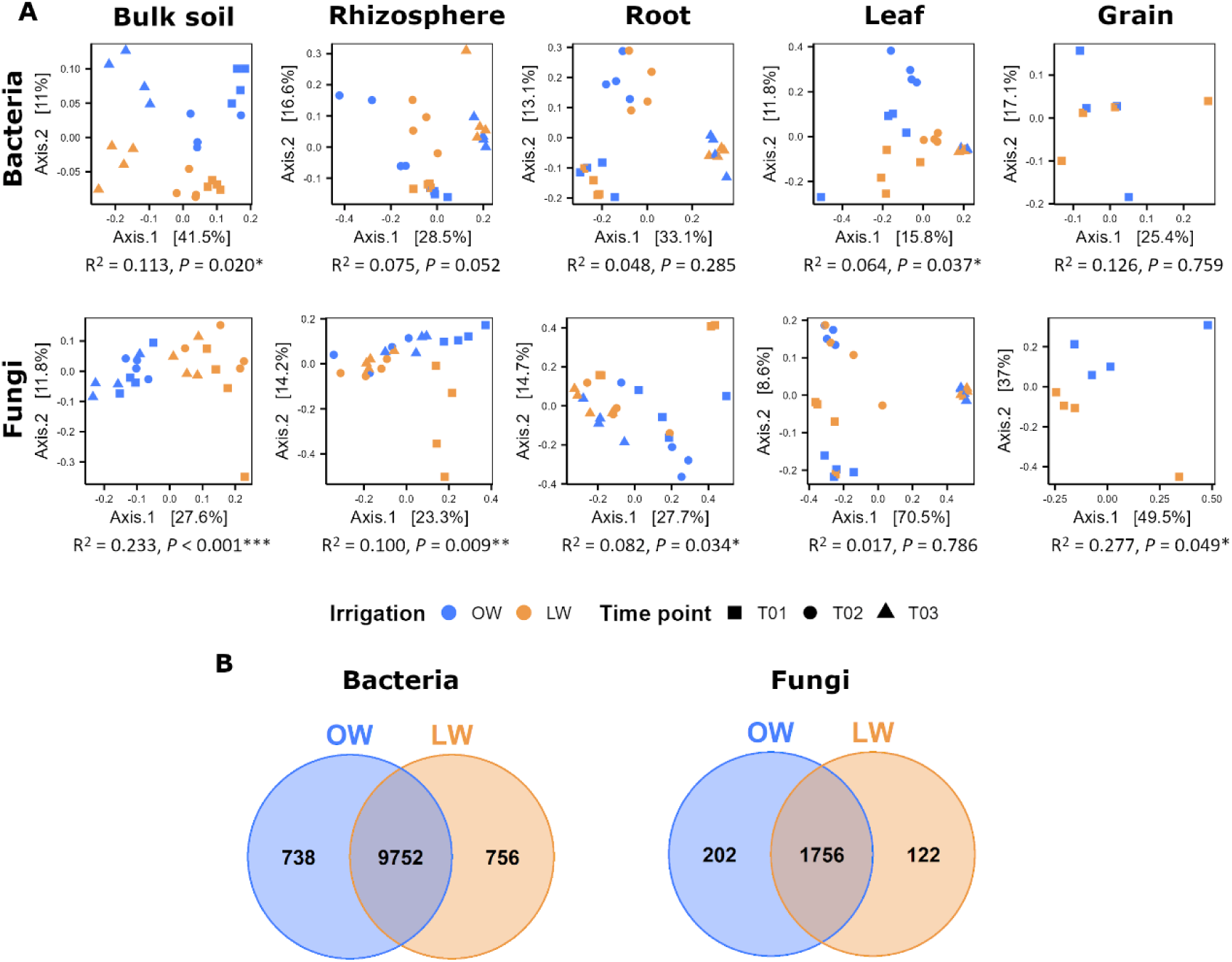
Impact of irrigation regime on maize-associated microbiome. A) Principal coordinate analysis (PCoA) based on Bray-Curtis dissimilarity of bacterial and fungal communities in each compartment unconstrained by irrigation regime (OW: optimal watering, LW: 30% watering reduction). The relative contribution (R^2^) of the factor “irrigation regime” on community dissimilarity and its significance was tested with PERMANOVA. Bulk soil, rhizosphere, root and leaf: n=24; grain: n = 8. B) Venn diagrams showing exclusive and shared bacterial and fungal ASVs between irrigation regimes.

We analysed the differences in abundance of the ASVs that were shared at least by two compartments by means of heatmaps and hierarchical clustering of the samples. Both in bacteria (Figure 5A) and in fungi (Figure 5B), an unequivocal separation between soil and plant niches was observed. Soil niche samples (bulk soil and rhizosphere) grouped first by irrigation regime and secondly by compartment, whereas plant niche samples grouped first by compartment and secondly by irrigation regime. This result suggest that the irrigation regime had a stronger impact on soil than on plant niche.

**Figure 5.**
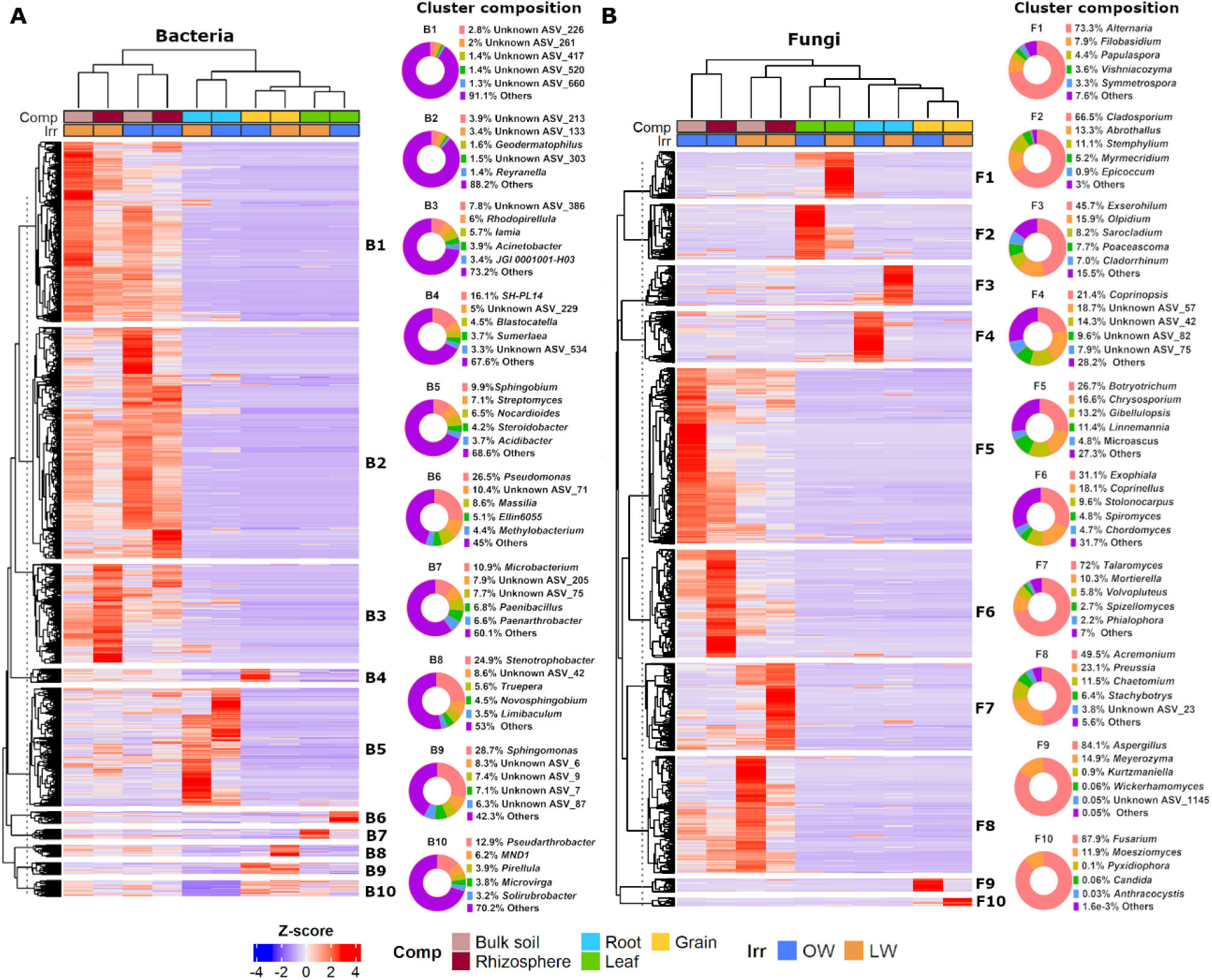
Differentially abundant clusters of bacterial and fungal ASVs from different maize-associated compartments under different irrigation regimes. Heatmap, hierarchical cluster and cluster composition diagram of (A) bacterial and (B) fungal ASVs shared by at least two compartments, divided by compartment and irrigation regime (OW: optimal watering, LW: 30% watering reduction). Cluster composition pie charts show the five most abundant genera within each cluster.

The hierarchical clustering of plant compartments showed clear differences between the bacterial and fungal assemblages. While in the bacterial assemblage leaves and grains grouped together and roots differentiated in a different branch (Figure 5A), it was the leaf that, in the case of fungi, differentiated from roots and grains (Figure 5B). This supports the results observed in the PCoAs of Figure 2 and highlights the different roles of roots and leaves in the assembly of bacterial and fungal microbiomes in the plant.

Heatmaps also revealed clusters of ASVs that were enriched in a specific compartment and irrigation regime (Figure 5, Supplementary Tables S7 and S8). This is especially remarkable in the case of fungi, where we can define a cluster specific for each compartment and irrigation regime combination (Figure 5B). Besides, each of these clusters had clearly predominant taxa, so we could define hallmark fungal genera for each compartment and irrigation regime (Supplementary Table S8). For example, clusters F1 and F2, dominated by ASVs of the genera *Alternaria* and *Cladosporium* respectively, showed groups of ASVs that were differentially enriched in the leaves, either at OW (cluster F2) or LW (cluster F1). Nonetheless, this situation is not so clear for bacteria (Figure 5A). Although some clusters showed ASVs enriched in a specific compartment and within an irrigation regime, such as clusters B4 and B8 in the grain or B6 and B7 in the leaf; others exhibited different patterns. For example, ASVs enriched in the roots of OW and LW clustered together in cluster B5, where *Sphingobium* and *Streptomyces* predominate, although presenting differences in abundance within the group. Another example is cluster B9, where the genus *Sphingomonas* was prevalent and enriched in the grain independently of the irrigation regime. Finally, cluster B10 showed a group of ASVs, predominated by *Pseudarthrobacter* and *MND1,* that were differentially depleted in the roots (Supplementary Table S7). It is important to note that the evenness in bacterial clusters is much higher than in fungal clusters (Table 2). Thus, while in the fungal clusters there are clearly prevalent ASVs, the frequencies of bacterial ASV are more similar and there are no clearly prevalent ones.

**Table 2.** Evenness of the bacterial and fungal clusters shown in Figure 5.

| Bacteria |  | Fungi |  |
| --- | --- | --- | --- |
| Cluster | Evenness | Cluster | Evenness |
| B1 | 0.87 | F1 | 0.42 |
| B2 | 0.84 | F2 | 0.55 |
| B3 | 0.79 | F3 | 0.54 |
| B4 | 0.90 | F4 | 0.63 |
| B5 | 0.79 | F5 | 0.54 |
| B6 | 0.79 | F6 | 0.54 |
| B7 | 0.84 | F7 | 0.44 |
| B8 | 0.85 | F8 | 0.38 |
| B9 | 0.80 | F9 | 0.35 |
| B10 | 0.88 | F10 | 0.25 |
| Average | 0.84 | Average | 0.46 |

### 3.5 Bacteria and fungi play different roles in increasing microbial network connectivity at LW

To explore the impact of irrigation regime in the interactions within the microbiome of each compartment, we conducted a co-occurrence network analysis (Supplementary Figure S6). Results showed clear differences in network parameters between OW and LW plants along the different compartments (Supplementary Table S9). Average degree was significantly higher in the bulk soil (*P* < 0.001) and root compartments (*P* < 0.001) of OW plants, and in rhizosphere (*P* < 0.001), leaves (*P* = 0.002) and grains (*P* < 0.001) assemblages of LW plants Positive edges (positive co-occurrence) were, in general, more frequent than negative edges (negative co-occurrence) (Figure 6B, Supplementary Table S9) especially for bacterial nodes. For fungal nodes, the number of positive and negative edges was more similar, and most of the negative edges corresponded to fungus-bacterium.

**Figure 6:**
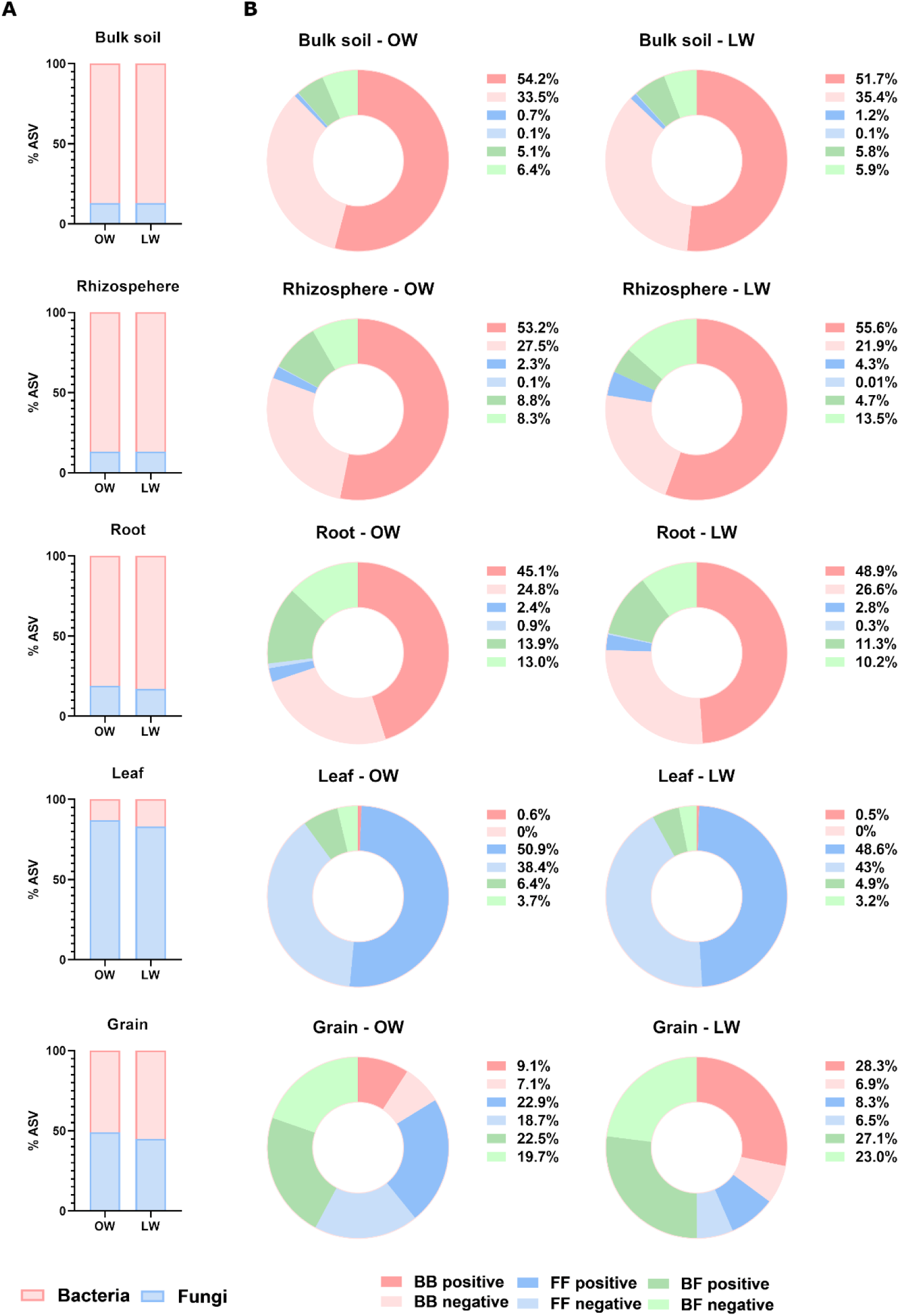
Intra- and interkingdom edges between bacterial and fungal taxa in co-occurrence networks in different maize-associated compartments under different irrigation regimes. A) Proportion of ASV of bacteria and fungi for each compartment and irrigation regime (OW: optimal watering, LW: 30% watering reduction). B) Pie charts representing the proportion of intra- and interkingdom edges between bacterial and fungal taxa in co-occurrence networks within each compartment and irrigation regime. BB represents bacteria-bacteria edges; FF represents fungi-fungi edges; BF represents bacteria-fungi edges.

There was a positive correlation between the number of ASVs in each sample and the number of nodes (*r* = 1, *P* < 0.001) and edges (*r* = 0.842, P = 0.002) (Figure 6A). Thus, in belowground compartments (bulk soil, rhizosphere, and root) bacterial ASVs were between 4 and 7 times more frequent than fungal ASVs, and more than 95% of the edges involved a bacterial node (Figure 6B). Conversely, fungal ASVs were prevalent in leaves by more than 5 times than bacterial ASVs, and 99% of the edges in this compartment involved fungi. In the grain, however, bacterial and fungal ASVs were equally prevalent, and the proportion of bacterial and fungal edges varied depending on the irrigation regime (Figure 6A and B). While edges connected mostly fungal taxa in OW, bacteria dominated the connections in LW. This difference was mostly due to fungus-fungus edges, which were decreased in LW plants, in earnings of positive bacterium-bacterium edges. Strikingly, the grain network at LW exhibited a 53.5% increase in the number of edges (OW:493; LW: 757), even though the number of nodes was similar (OW: 145; LW: 144) in both irrigation regimes. This could be explained by a 2.2-fold increase in the number of edges, mostly positive, in the bacterial nodes at LW in comparison to OW.

Further, we defined “network hubs” as nodes with high values of degree and betweenness centrality (≥ 𝑃_90_, Supplementary Figure S7). In total, 319 bacterial ASVs and 47 fungal ASVs were identified as hubs (Supplementary Table S10). The number of bacterial hubs correlated with the number of analysed ASVs (*r* = 0.89, *P* < 0.001), but there was no correlation between the number of analysed ASVs and the number of fungal hubs (*r* = -0.03, *P* = 0.924). For both bacteria and fungi, the rhizosphere was enriched in hubs in comparison, for example, to bulk soil which had similar number of nodes (in fact, there were no fungal hubs in the bulk soil). Interestingly, the number of fungal hubs in LW was 2.6-fold higher than in OW (OW: 14; LW: 36), whereas in the case of bacteria the numbers were similar for both irrigation regimes (OW: 191; LW: 195). This was mainly due to an enrichment in the number of fungal hubs in the rhizosphere at LW, which was 4.4-fold greater than in OW (OW: 7 hubs, LW: 31 hubs). Most of the edges of these fungal hubs at LW were due to negative co-occurrence with bacteria (Supplementary Table S11), while in OW most of the connections were due to positive co-occurrence with bacteria.

The rise in the number of bacterial positive edges exhibited in the grain at LW indicates that bacteria may play a relevant cooperative role in the establishment of grain microbial networks under those conditions. On the other hand, fungi seemed to have a key competitive function in the rhizosphere, since fungal hubs were enriched in comparison to OW, and these hubs had predominant negative edges (negative co-occurrence) with bacteria. Fungi must also have an important role in the establishment of the foliar microbiome and its co-occurrence networks. Thus, our results suggest that bacteria and fungi play differential roles in the conformation of microbial networks in different compartments at LW.

## 4. Discussion

During the last decade, a growing number of studies describe the plant associated microbiomes (mostly root associated bacteria) of different host species and in different environments. However, it is still necessary to expand the body of data under different conditions to extract robust conclusions about the basic principles of the plant microbiome conformation (Burz et al., 2023). This knowledge is essential for the establishment of models that enable a better understanding of microbiome functioning and their management in the agroecosystems (Compant et al., 2019; Burz et al., 2023).

With this study, we provide a comprehensive analysis of the structure of both bacterial and fungal communities in the bulk soil, rhizosphere, roots, leaves and grains of open-field cultivated maize plants during different stages of the crop cycle under two different irrigation regimes. Our results highlight the dynamic nature of the microbial communities and the contrasting patterns and interconnections between bacterial and fungal assemblages, evidencing that single time points and kingdom exclusive studies provide insufficient information on the complexity of the plant microbiome.

In this work we take advantage of the irrigation needs of maize crop in a semi-arid agricultural area to study the effect of an environmental stressor (water scarcity) without changing other conditions in an open-field trial. Our results show that plant microbiome conformation is a very robust process and that only minor changes, which mainly affect fungi, can be associated to the applied water stress. Indeed, bacterial and fungal assemblages show different diversity dynamics that point to kingdom-specific selective processes during maize microbiome conformation. Based on our findings and previous literature, we discuss below the different factors that may be affecting this process (Figure 7).

**Figure 7.**
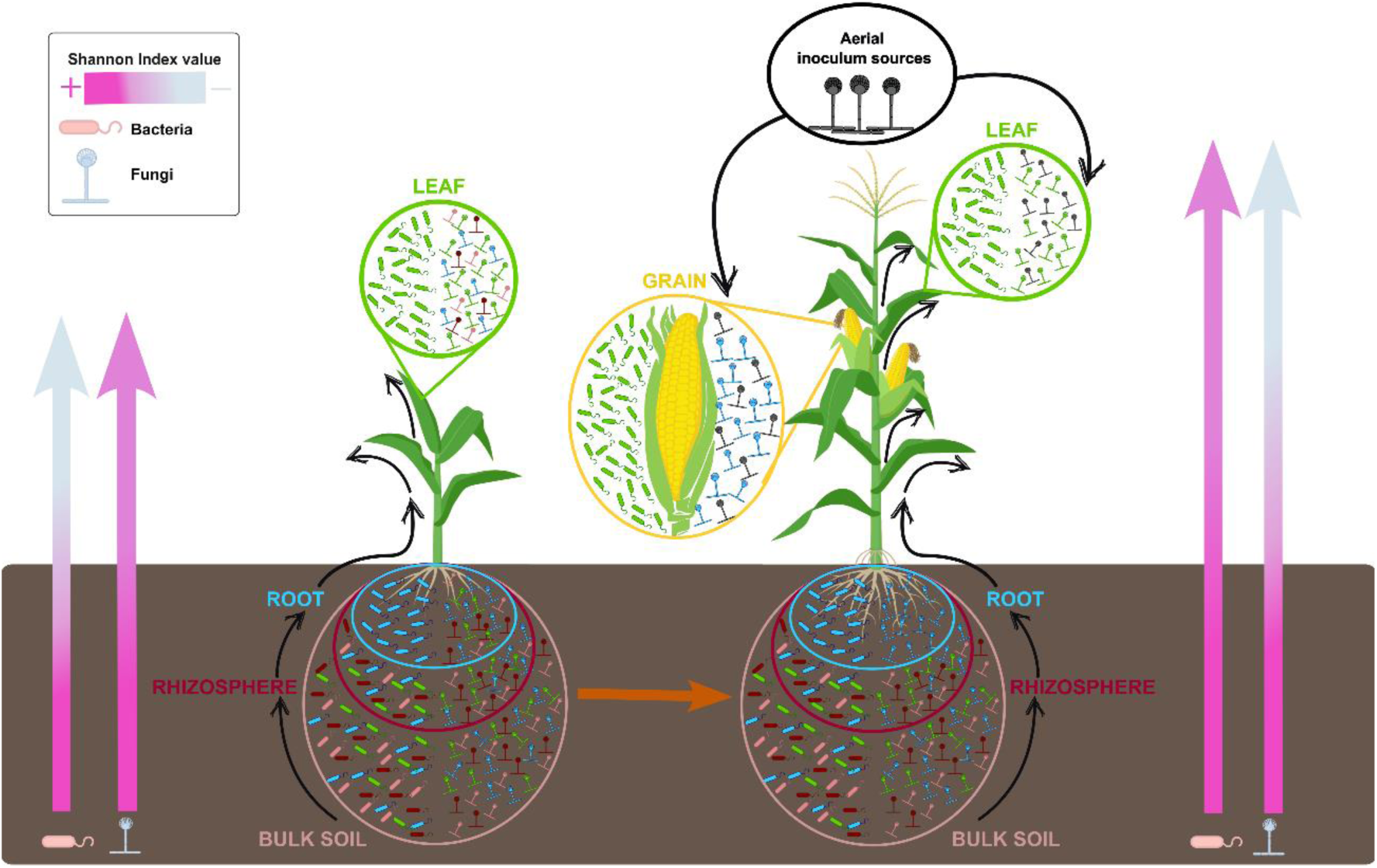
A model depicting maize bacterial and fungal microbiome conformation. Plant selects bacteria and fungi from soil compartments. This host effect is stronger in bacteria than in fungi since the bacterial microbiomes of the roots and the leaves are differentiated since the beginning of plant development. On the contrary, the plant needs more time to shape the fungal community, which is more similar to soil compartments at earlier time points and tend to differentiate with plant development. Maize grain bacterial communities are closer to those in the leaf, while fungal communities are more similar to those in the root. Fungal communities of the aboveground organs (leaf and grain) have a relevant proportion of exclusive taxa whose inoculum may have an aerial origin. Bacterial alpha-diversity (Shannon index) in plant compartments is strongly reduced at the beginning of the plant cycle, while fungal diversity decreases towards the end of the plant cycle due to the prevalence of few taxa.

### 4.1. Plant selection of soil microbiota prevails over environmental effects

Our work indicates that maize microbiota is mainly derived from soil and gradually filtered by different compartment niches (Figure 3A, Supplementary Figure S5 and Supplementary Table S5), in consistence with previous findings (Tkacz et al., 2020; Xiong et al., 2021a; Zhang et al., 2022). The strong relatedness between soil and root-associated microbiota has been demonstrated by numerous studies (Lundberg et al., 2012; Zarraonaindia et al., 2015; Liu et al., 2019; Thiergart et al., 2019). Environmental parameters and soil properties have been pointed out as determining factors of soil and root microbial community composition, affecting differentially both bacterial and fungal communities (Bahram et al., 2018; Thiergart et al., 2019). Overall, the bacterial microbiomes seem to be mainly impacted by edaphic factors, such as pH, whereas climatic conditions determine fungal community profiles. In this line, our results also show that an environmental stressor like water limitation had a stronger influence on the microbiome structure of fungi than bacteria in the belowground compartments (Figure 4A and Supplementary Table S4). Our clustering analysis also shows a strong effect of irrigation regime on soil and rhizosphere compartments (Figure 5). However, the plant compartment prevailed over the irrigation regime in the clustering of the microbial assemblies within the plant. Actually, although location, soil or climate define the ensemble of taxa from which the plant can harbor its microbiome, it is already rather clear that the host plant determines the microbes that finally reach the inner tissues of the plants (Robbins et al., 2018; Wang et al., 2019; Xiong et al., 2021a, 2021b; Zhang et al., 2022).

Our results show that the rhizosphere shares most of its microbiome with the rest of the compartments (Figure 3 and Supplementary Table S5). Indeed, because of the intense biochemical activity found in the vicinity of the roots, the rhizosphere is the first layer of soil on which the plant host can influence the conformation of its microbiome. In the rhizosphere, root exudates play different roles that contribute to the fine-tuning of the root microbiome (Hu et al., 2018; Huang et al., 2019; Stassen et al., 2021). On the one hand, they can act as attracting signals or as carbon substrates for microbial growth (Bulgarelli et al., 2013; Sasse et al., 2018; Huang et al., 2019). On the other hand, the secretion of plant specialized metabolites, for example those with antimicrobial activity, also determines the microbiota that reaches the roots due to the different levels of tolerance or susceptibility to these compounds (Huang et al., 2019; Thoenen et al., 2023). In maize, benzoxazinoids (BXs) are a good example of this dual and selective function. For instance, it has been reported that wild-type maize producing the BX compound DIMBOA promote the colonization of maize roots by the plant-beneficial bacteria *P. putida* KT2440 in comparison to DIMBOA-deficient *bx1* mutant (Neal et al., 2012). Recently it has also been shown that tolerance to BXs strongly determines the composition and structure of maize root bacterial communities (Thoenen et al., 2023). It is yet to be elucidated if different plant selective mechanisms affect to bacterial and fungal communities, and which are the factors that explain the differential recruitment across microbial kingdoms, since as our study highlights, the processes of microbiome establishment are different for bacteria and fungi.

### 4.2. Bacterial diversity in plant compartments is strongly reduced at the beginning of the plant cycle, while fungal diversity decreases towards the end of the plant cycle due to the prevalence of few taxa that are affected by irrigation regime

Previous studies unveiled that compartment and developmental stage are key factors shaping microbial communities of plants (Cregger et al., 2018; Xiong et al., 2021a; Zhang et al., 2022). Here, we also show that soil and plant compartments are the main drivers of maize associated microbial communities during plant development (Supplementary Figure S4, Supplementary Table S4), although with some differences in bacterial and fungal assemblages. Both bacteria and fungi exhibited an abrupt reduction in richness in plant compartments compared to soil compartments (Figure 1A). Bacterial assemblages however, with ten times higher richness than fungi in soil compartments, showed a stronger reduction in aboveground compartments than fungal assemblages. This confirms the results of Xiong et al., (2021a) and supports the hypothesis of a stronger plant effect in the recruitment of the bacterial than the fungal microbiome. The severe effect on richness reduction in the plant remained stable during the whole plant cycle. However, time did modulate microbial Shannon index, with different effects in bacteria and fungi. While Shannon index of bacteria in the different compartments tended to be stable or even increase towards the end of the cycle, fungal communities exhibited a staggered decrease in this parameter from bottom (soil) to upper compartments in the plant (leaf) (Figure 1B and 7), reflecting a decrease in evenness due to the prevalence of few taxa (Supplementary Figure S3). Although we have not found previous works that monitor alpha-diversity, either Shannon index or evenness, of bacteria and fungi along the soil-plant continuum, some studies have revealed comparable results in specific plant compartments at different time points. For example, Bourceret et al. (2022) also showed that fungal Shannon index in maize roots fell from the vegetative to the reproductive stage, while bacterial remained stable. Similar results were reported by another study monitoring leaf microbiome dynamics throughout the natural growing season of *A. thaliana* (Almario et al., 2022). Although these two studies did not specifically look to the changes in evenness, they both found an increase in the relative abundance of certain fungal OTUS with time that could be reducing evenness and, thus, fungal alpha-diversity.

Besides, we found a compartment-kingdom specific differentiation in beta-diversity that was enhanced during plant cycle. Our analyses showed a gradual differentiation of bacterial assemblages at earlier time points that changed towards the end of the cycle, with the roots showing a strong differentiation from the rest of the compartments (Figure 2). The fungal communities of the different compartments tended to overlap at earlier time points but showed a strong differentiation at T03, with an increase in compartment exclusive ASVs towards the plant compartments that was intensified in the leaf (Figures 2 and 3, Supplementary Figure S5). The substantial reduction in alpha-diversity in plant compared to soil compartments, along with the increase in assemblage differentiation with time, support the idea of a dynamic compartment-based selection process driven by the plant, with kingdom-specific mechanisms of recruitment. This process would have led to the diversification of bacterial communities from the soil to the different plant niches, in contrast with the prevalence of fungal exclusive taxa in each plant compartment, strongly affected by the irrigation regime (Figure 5).

### 4.3. Bacterial communities differentiate mostly in the roots at the end of maize cycle, and fungal communities differentiate in the leaves with a probable contribution of aerial inoculum

The large proportion of shared taxa between the roots and the leaves reported by some studies suggest that, after successful colonization, microorganisms might be transmitted from the roots to the phyllosphere through the xylem or nonvascular tissue (Bai et al., 2015; Ibekwe et al., 2020; Tkacz et al., 2020; Xiong et al., 2021a, 2021b; Zhang et al., 2022). The fact shown in our study that most of the microbial taxa were shared among the different soil and plant compartments (Figure 3A and D) and the presence of a core microbiota during plant cycle (Figure 3B and E) further supports this idea. Nonetheless, the pressure of the immune system and the presence of specific metabolites or nutrients exert a selection that defines the ensemble of microorganisms of different plant microhabitats or compartments, such as the roots or the leaves (Hassani et al., 2019; Fitzpatrick et al., 2020; Trivedi et al., 2020; Mesny et al., 2023). Our results showed a strong differentiation of the bacterial assemblage in the roots and the fungal assemblage in the leaves at the end of the cycle (Figure 2). The structural and physiological differences between the roots and leaves along with the exposure to soil and air, respectively, might explain the differential microbial composition of these compartments (Müller et al., 2016; Fitzpatrick et al., 2020). Besides the predominance of microbes from the soil, our data also suggest that additional aerial sources of inoculum in the aboveground organs might be especially relevant for fungi (Figure 3, Supplementary Figure S5). Bell-Dereske and Evans (2021) also showed that aerial transmission of fungi is an important driver of the leaf endophytic fungal community of switchgrass (*Panicum virgatum*). Another study that compared the phylloplane of plastic plants with that of maize leaves revealed a greater proportion of microorganisms originated from the air in the fungal than in the bacterial assemblage of the leaves (Xiong et al., 2021a). Thus, besides the internal transmission of soil microorganisms along the plant, it is necessary to take into account the aerial inoculum as external contribution to the fungal microbiome of maize aboveground organs. This external contribution entails further opportunities for the establishment of taxa different from the ones present in the soil.

### 4.4. Maize grain bacterial communities are closer to those in the leaf, while fungal communities are more similar to those in the root

The seed microbiota can be considered both an endpoint for microbial community assembly and also, together with soil, an additional starting point for the microbiome of the new plant (Shade et al., 2017; Rochefort et al., 2021). It has been suggested that the transference of microorganisms from other plant organs through the xylem and non-vascular tissues to the seeds represents one of the main processes of seed microbiome assembly and vertical transmission to the next generation of plants (Shade et al., 2017; Nelson, 2018). However, it is difficult to find studies focused on the relative contribution of the microbiomes of other plant compartments, such as the roots or the leaves, on seed microbial assemblages during its development in the mother plant. Our analysis showed that grain-associated bacterial and fungal assemblages grouped differently, being bacterial communities closer to those of leaf samples and fungal assemblages more similar to those in the root (Figure 2, 5 and 7). This suggests that the bacterial community of the grain mainly derived from the leaves, while the fungal community originated from the roots. Xiong et al. (2021a) also showed a highest contribution of the leaves for the bacterial assemblage and the roots for the fungal assemblage of maize grains. Since it has been shown that vertically transmitted seed microbiome can improve germination, plant survival and productivity (Barret et al., 2016; Shade et al., 2017; Nelson, 2018; War et al., 2023), it is important to increase our understanding of seed microbiome assembly and the origin of vertically transmitted bacteria and fungi. This may allow us to engineer microbiomes to avoid plant pre- and post-harvest diseases and increase crop productivity (Shade et al., 2017; Orozco-Mosqueda et al., 2018; Arif et al., 2020).

### 4.5. Fungi play leading roles in the maize microbiome shifts driven by water stress

Maize is a high water-demanding crop, which means that potential reductions in water availability, either due to irrigation water restrictions or drought periods, substantially impact maize productivity (Daryanto et al., 2016; Leng and Hall, 2019; Sah et al., 2020). In this work, we showed that a reduction of 30% of the optimal irrigation of maize grown in an open field causes a water stress in the plant that leads to a significant reduction of chlorophyll index, shoot biomass and yield (Table 1). Albeit the observed negative effect on plant growth and yield, the microbiota of LW plants did not show an overall significant shift. However, we showed compartment-specific variations, which mainly affected the fungal community (Supplementary Table S3, Figure 4A). Our heatmap analysis revealed clear irrigation-specific clusters of fungal ASVs that were differentially abundant for each compartment (Figure 5B), suggesting that irrigation-specific restructuring is occurring within each compartment in the fungal assemblage. These clusters were dominated by specific taxa that might play key roles in the adjustments of the microbial community under water stress. These results can be related to the previous findings indicating that fungal communities could be more influenced by atmospheric environmental factors, such as rainfall or air temperature, while soil characteristics and plant genotypes might be relevant determining factors of bacterial communities (García et al., 2013; Coleman-Derr et al., 2016; Bahram et al., 2018; Thiergart et al., 2019; Bell-Dereske and Evans, 2021).

Our analysis showed compartment-specific variations in key topological properties of microbial networks in response to water stress. Average degree was significantly higher in the rhizosphere, leaves and grains assemblages of LW plants. Moreover, the microbial networks of the grain at LW showed a 53.5% increase in the number of edges, mostly due to positive co-occurrence among bacterial taxa. Increases in network complexity are consistent with previous studies that suggest that drought periods and water stress could enhance the connectivity of microbial networks as a “cry for help”, so plants can recruit specific microbes to alleviate environmental stresses (de Vries et al., 2018; Rodriguez et al., 2019; Liu et al., 2020; Yuan et al., 2021; Zhai et al., 2024). Our results also revealed an increase in the negative co-occurrence between fungal and bacterial taxa in the rhizosphere of LW plants. This was mainly due to a rise in the number of negative edges of fungal nodes, with an enrichment in fungal hubs. Other studies have reported that negative bacteria-fungi edges in the rhizosphere become more intense under water stress (Bazany et al., 2022), suggesting that this behaviour could be due to the strong competition for nutrients, mainly from root exudates (Durán et al., 2018). We also observed an increase in negative fungus-fungus correlations in the leaf in LW compared to OW. Indeed, positive and negative correlations in the co-occurrence of different taxa represent co-oscillation and competition in the ecosystem, respectively (Coyte et al., 2015). Positive correlations among microbial taxa can stand for different cooperative interactions, essentially cross-feeding, or just indicate that they are responding in tandem to an environmental disturbance (Mesny et al., 2023). Because of this dependence link, modelling studies predict that increased positive co-occurrence could negatively influence network stability. On the other hand, negative co-occurrence tends to balance the microbiome and increases its stability under the exposition to stressors (Coyte et al., 2015; de Vries et al., 2018). Therefore, the increase in negative co-occurrence might indicate a way of reaching network stability in specific compartments in which fungi must play an essential role. A recent study on tobacco leaves revealed that fungal networks were more complex than bacterial networks, with strong negative correlations between fungi and bacteria (Zhou et al., 2021). Thus, our results add evidence to the significant role of fungi on improving microbial community stability, pointing to their possible relevance in increasing resilience to water stress. Overall, we demonstrate that not only compositional shifts but also network complexity and intra- and inter-kingdom connections mediate the response of maize plant microbiomes to the water stress due to irrigation restrictions, highlighting the strong need of conducting multi-kingdom analyses to better understand how microbial ecosystems function.

## 5. Conclusions

We provide a model that describes plant microbiome conformation by studying maize associated bacterial and fungal communities in the different plant compartments during plant cycle and at different irrigation regimes, integrating other published results. Our work confirms that the assembly of the plant microbiome is a robust compartment-based dynamic process that evolves with plant development. We showed that water stress does not have a severe impact on maize microbial diversity, although relevant changes can be accounted for within specific compartments that affect differently to bacteria and fungi. Whether the observed impacts contribute to plant performance under stress still needs to be further investigated. Our results also suggest that the mechanisms of recruitment and assembly may be different for bacteria and fungi, with members from different kingdoms playing different roles in plant functioning and response to environmental stressors. Thus, our study highlights the importance of conducting multikingdom analyses that contribute to a holistic understanding of the dynamics and evolution of the microbial assemblages along the whole plant, providing data that can be used for further studies. A deep comprehension of plant-microbiome-environment interactions is essential for the development of sustainable solutions which enable the manipulation of plant and seed-associated microbiomes to enhance plant resilience and productivity under the current and future climatic scenarios.

## Supporting information

Supplementary table 1_Reads and read length

Supplementary table 2_Phyla, genus and ASV counts

Supplementary table 3_Statistics alpha diversity

Supplementary table 4_Permanova

Supplementary table 5_Venn frequencies and percentages

Supplementary table 6_Venn_taxa

Supplementary table 7_heatmap_cluster_genus_16S

Supplementary table 8_heatmap_cluster_genus_ITS

Supplementary Table 9_topological properties networks

Supplementary Table 10_Hub nodes taxonomy

Supplementary Table 11_Hub nodes co occurrences

## Additional Files

**Additional File 1: Supplementary Methods**

“Additional File 1_Supplementary methods.pdf”

**Additional File 2: Supplementary Figures**

“Additional File 2_Supplementary Figures.pdf”

**Additional File 3: Supplementary Table S1:** Total number of reads and read length before and after quality check.

“Supplementary table 1_Reads and read length.xlsx”

**Additional File 4: Supplementary Table S2:** Assigned numbers of bacterial and fungal phyla, genera and ASVs at different compartments, time points and irrigation regimes.

“Supplementary table 2_Phyla, genus and ASV counts.xlsx”

**Additional File 5: Supplementary Table S3:** Alpha-diversity indicated by Chao1 and Shannon under compartments, time points and irrigation variations with related statistical analysis.

“Supplementary table 3_Statistics alpha diversity.xlsx”

**Additional File 6: Supplementary Table S4:** PERMANOVA analysis under different compartments, time points and irrigation regimes.

“Supplementary table 4_Permanova.xlsx”

**Additional File 7: Supplementary Table S5:** Frequencies and percentages of shared and exclusive bacterial and fungal ASVs under different compartments, time points and irrigation regimes.

“Supplementary table 5_Venn frequencies and percentages.xlsx”

**Additional File 8: Supplementary Table S6:** Taxonomic composition of the bacterial and fungal groups derived from Venn diagrams under different compartment, time points and irrigation regimes.

“Supplementary table 6_Venn_taxa.xlsx”

**Additional File 9: Supplementary Table S7:** Taxonomic composition of the bacterial clusters derived from hierarchical clustering at the genus level.

“Supplementary table 7_heatmap_cluster_genus_16S.xlsx”

**Additional File 10: Supplementary Table S8:** Taxonomic composition of the fungal clusters derived from hierarchical clustering at the genus level.

“Supplementary table 8_heatmap_cluster_genus_ITS.xlsx”

**Additional File 11: Supplementary Table S9:** Major topological properties of co-occurrence networks of each compartment and irrigation regime.

“Supplementary Table 9_topological properties networks.xlsx”

**Additional File 12: Supplementary Table S10:** List of network hubs with taxonomy, betweenness centrality and degree.

“Supplementary Table 10_Hub nodes taxonomy.xlsx”

**Additional File 13: Supplementary Table 11:** Proportion of positive and negative intra- and interkingdom edges of hub nodes in co-occurrence networks within each compartment and irrigation regime.

“Supplementary Table 11_Hub nodes co occurrences.xlsx”

## Authors’ contributions

S.D.-G., S.G.-B., C.G.-S., P.M., F.B. and S.S conceptualized and designed the research. S.D.-G. and P.M. collected and processed the samples. S.G.-B. processed the raw sequences. S.D.-G. and S.G.-B. conducted the downstream bioinformatic analysis. S.D.-G. prepared the tables and figures. S.D.-G., S.G.-B., C.G.-S. and S.S contributed to the text of the main manuscript. All authors have read and agreed to the published version of the manuscript.

