## Additional File 1_Supplementary methods for "Maize associated bacterial and fungal microbiomes show contrasting conformation patterns that are resilient to water availability"

**1. Irrigation program for open-field maize plants subject to optimal (OW) and limited (LW) watering condition.**

| | CROP ESTABLISHMENT | VEGETATIVE DEVELOPMENT | | REPRODUCTIVE STAGE | SENESCENCE | | |
|---|---|---|---|---|---|---|---|
| | | Stage 1 | Stage 2 | | Stage 1 | Stage 2 | Stage 3 |
| No. days per period | 20 | 17 | 18 | 40 | 10 | 10 | 10 |
| Watering regime until | 28/05/18 | 14/06/18 | 02/07/18 | 11/08/18 | 21/08/18 | 31/08/18 | 10/09/18 |
| Kc[1] | 0.4 | 0.79 | 1.20 | 1.20 | 0.92 | 0.63 | 0.35 |
| Eto (mm/day) | 4.80 | 5.25 | 5.72 | 5.55 | 5.16 | 4.73 | 4.32 |
| Rainfall (mm/day) | 0.30 | 0.14 | 0.20 | 0.10 | 0.29 | 0.35 | 0.62 |
| Effective rainfall (mm/day) | 0.1 | 0.0 | 0.0 | 0.0 | 0.1 | 0.1 | 0.3 |
| Etc (mm/day) | 1.92 | 4.14 | 6.86 | 6.66 | 4.73 | 2.99 | 1.51 |
| GIWR (mm/day) | 1.82 | 4.14 | 6.84 | 6.66 | 4.64 | 2.86 | 1.17 |
| NIWR (mm/day) | 1.80 | 4.10 | 6.77 | 6.60 | 4.59 | 2.83 | 1.16 |
| TIWR (mm/day) | 2.37 | 5.39 | 8.91 | 8.68 | 6.04 | 3.72 | 1.53 |
| Watering time (h/day) | 0.51 | 1.16 | 1.91 | 1.86 | 1.29 | 0.80 | 0.33 |
| Watering time in OW (min/day) | 30 | 69 | 115 | 112 | 78 | 48 | 20 |
| Watering time in LW (min/day) | 21 | 49 | 80 | 78 | 54 | 33 | 14 |

| | | | |
|---|---|---|---|
| Eto | Reference evapotranspiration | Kl | Location correction coefficient |
| ETC | Reference crop evapotranspiration | Kr | Climatic variation correction coefficient |
| GIWR | Gross irrigation water requirements | Ka | Advection correction coefficient |
| NIWR | Net irrigation water requirements | Ae | Application efficiency |
| TIWR | Total irrigation water requirements | CU | Uniformity coefficient |

Calculations were made on long-term data from the closest climate station, located in San Javier (37° 47' 20'' N, 0° 48' 12'' O; IMIDA 2018).

[1]Corn Kc was adapted from CROPWAT (version 8.0) and seed breeder information.

### 2. Processing of soil and plant material until DNA extraction

BULK SOIL COMPARTMENT: Soil material from the top 30 cm surrounding layer of each plant was collected in 2 mL Eppendorf tube with lysis matrix from FastDNA™ Spin Kit for Soil (MP Biomedicals).

RHIZOSPHERE COMPARTMENT: Roots were separated from the rest of the plant and shaken to remove the excess of soil. Roots were washed twice in two 500 mL plastic recipients with 100 mL of sterile distilled water to remove attached soil particles. The 200 mL of washing water (containing soil particles) from the same plant, were mixed and 30 mL were transferred to a 50 mL falcon. Falcon tubes were centrifuged at 4,000xg for 15 min. After centrifugation, 95% of supernatant was discarded. Pellet was resuspended into the remaining supernatant. Using a cut P1000 tip, 300 µL of the mixture were transferred to a 2 mL tube with lysis matrix from FastDNA™ Spin Kit for Soil (MP Biomedicals).

ROOT COMPARTMENT: Roots were gently washed with tap water in order to remove any remaining soil residue. Roots were cut into small sections with pruning shears, which were disinfected with a 20% bleach solution between samples. Root sections were wrapped in aluminum foil for their storage.

LEAF COMPARTMENT: Four circles of ~1 cm of diameter were collected from the youngest unfolded leaf of each plant and introduced in a 2 mL Eppendorf tube filled with sterile glass beads (2.7 mm Ø, Carl Roth GmbH & Co.).

GRAIN COMPARTMENT: Grain samples were only collected for T03. Mature cobs from the same plant were manually dekerneled, grains were pooled and transferred to 15 mL falcon.

All samples were kept at -80ºC until DNA extraction.

Bulk soil, rhizosphere, and leaf material were homogenized using FastPrep-24™ 5G homogenizer (MP Biomedicals), while roots and grain samples were disrupted with a mortar and liquid nitrogen.
