## Additional File 2_Supplementary Figures for "Maize associated bacterial and fungal microbiomes show contrasting conformation patterns that are resilient to water availability"

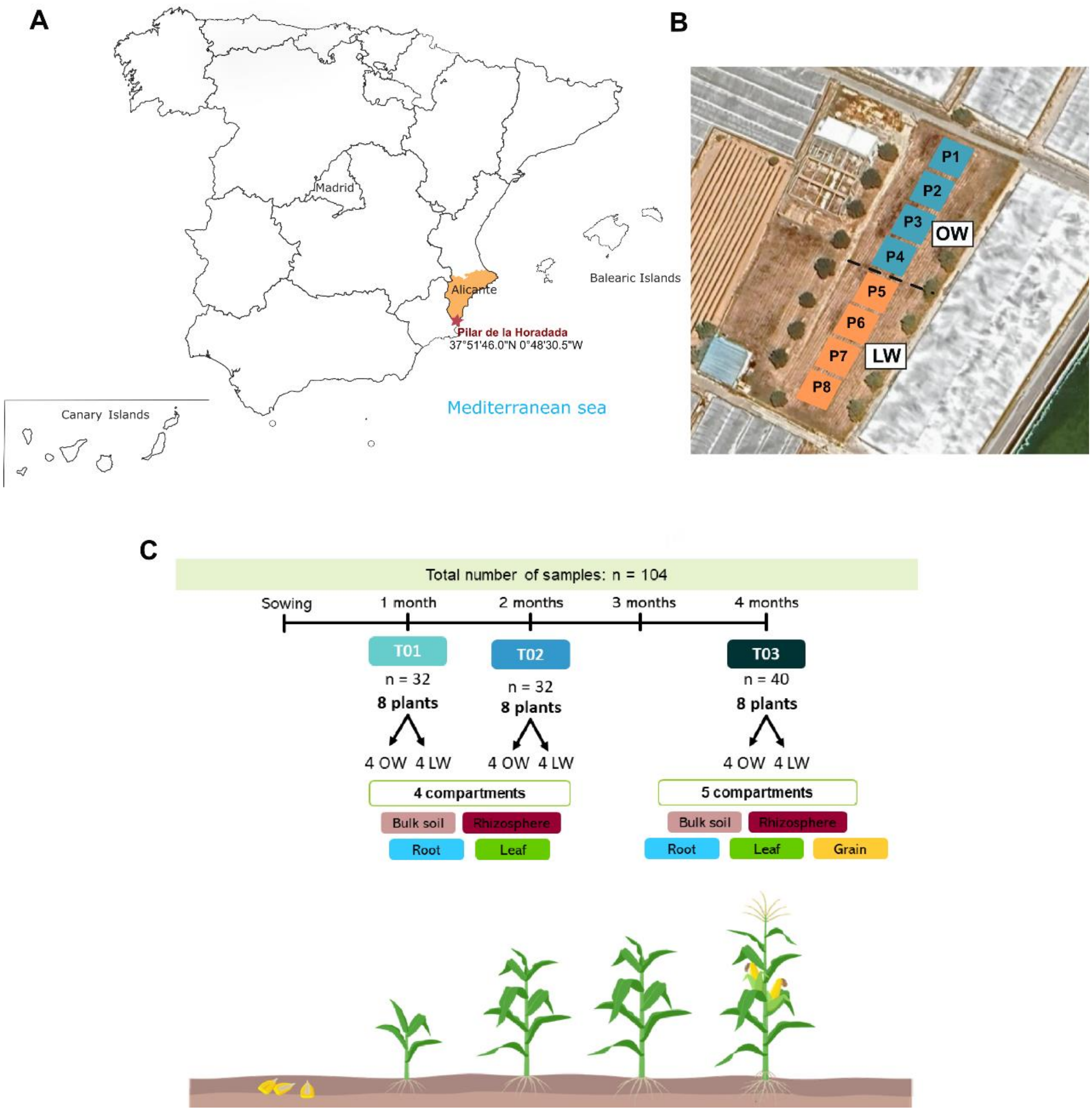


**Supplementary Figure S1:** A) Geographical location of the experimental station. B) Plot distribution in the experimental field. C) Sampling scheme. OW: optimal watering, LW: 30% watering reduction.

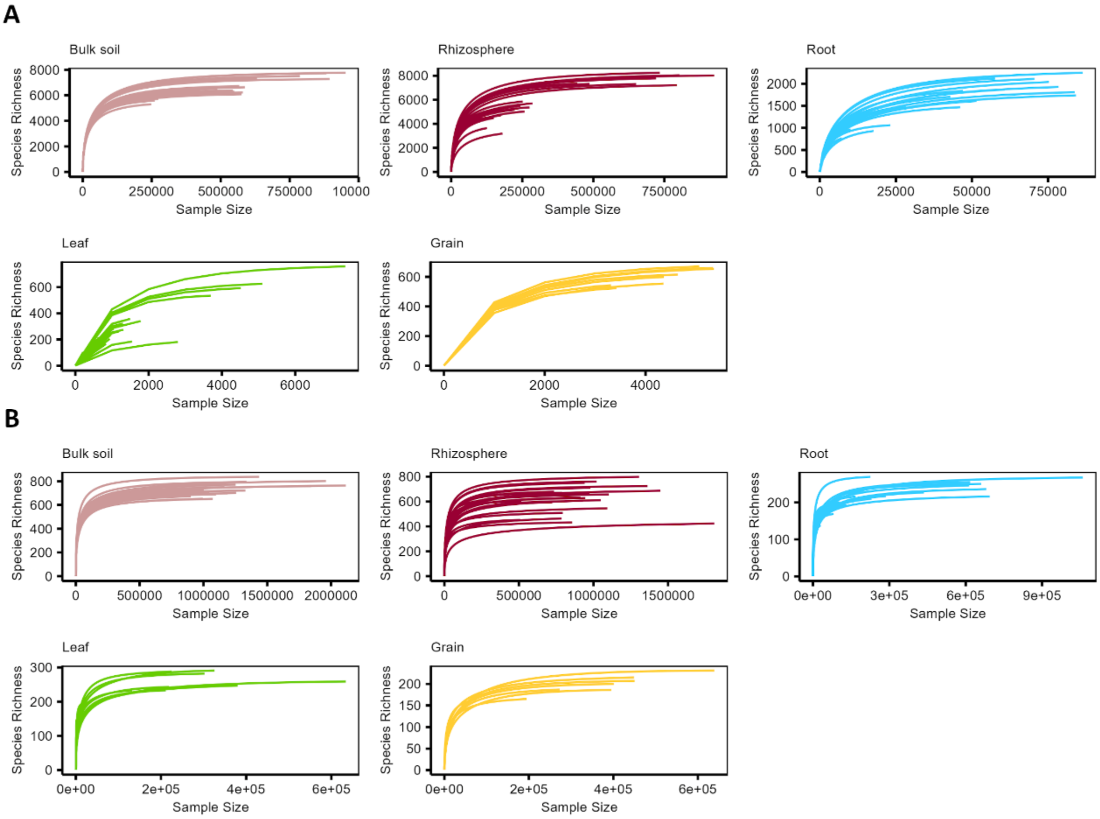


**Supplementary Figure S2:** Rarefraction curves of the bacterial (A) and fungal (B) datasets divided by compartment.

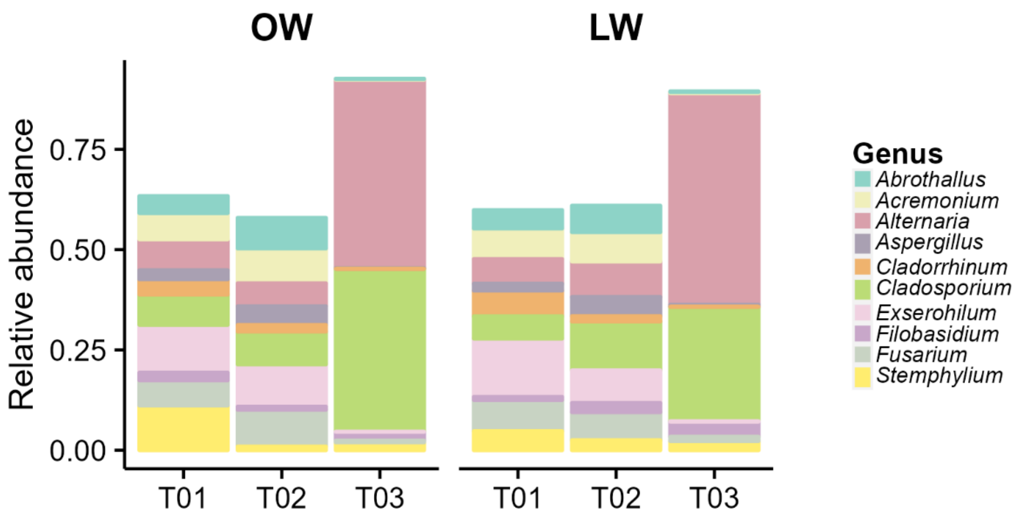


**Supplementary Figure S3:** Top 10 fungal genera of leaf microbiome at different time points (one -T01-, two -T02- and four -T03- months after sowing) and irrigation regimes (OW: optimal watering, LW: 30% watering reduction)

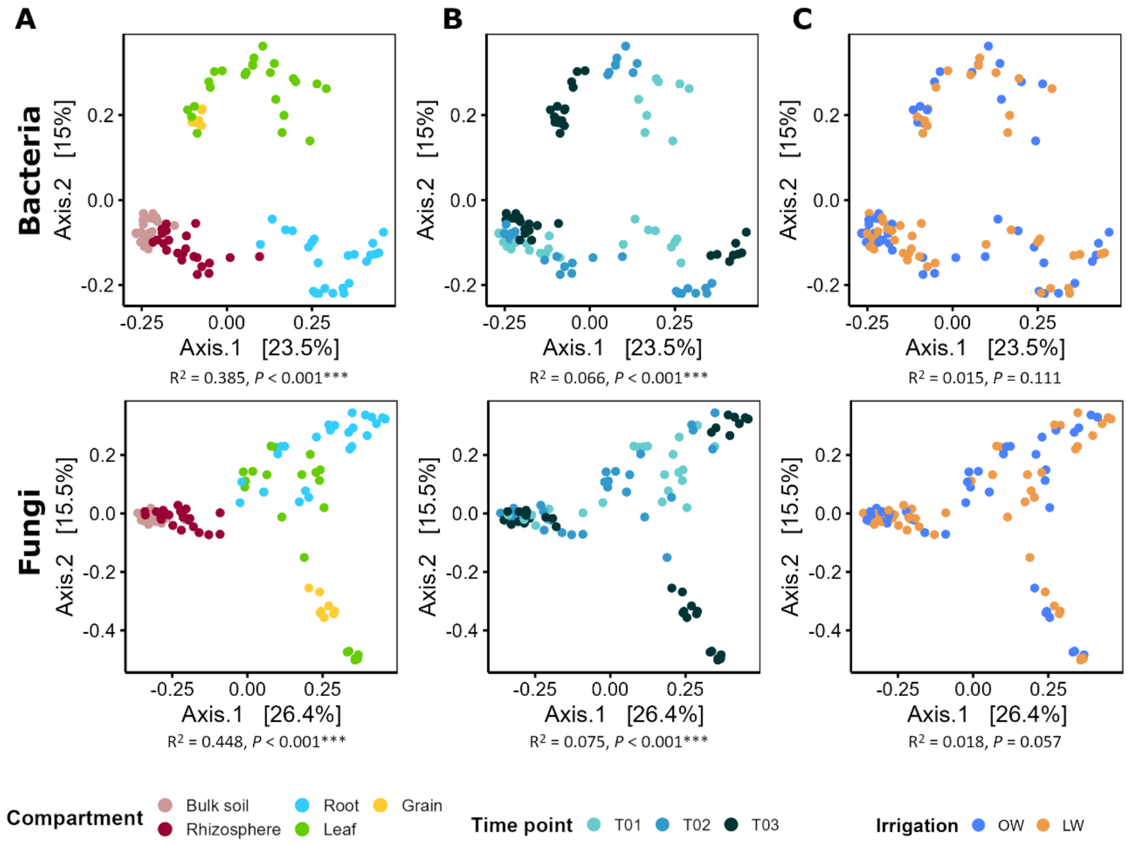


**Supplementary Figure S4: General beta-diversity patterns of maize-associated microbiota.** Principal coordinate analysis (PCoA) based on Bray-Curtis dissimilarity depicting the distribution patterns of bacterial and fungal communities unconstrained by (A) compartment, (B) time point (one -T01-, two -T02- and four -T03- months after sowing) and (C) irrigation regime (OW: optimal watering, LW: 30% watering reduction) (n = 104). The relative contribution ($R^2$) of different factors on community dissimilarity and its significance was tested with PERMANOVA.

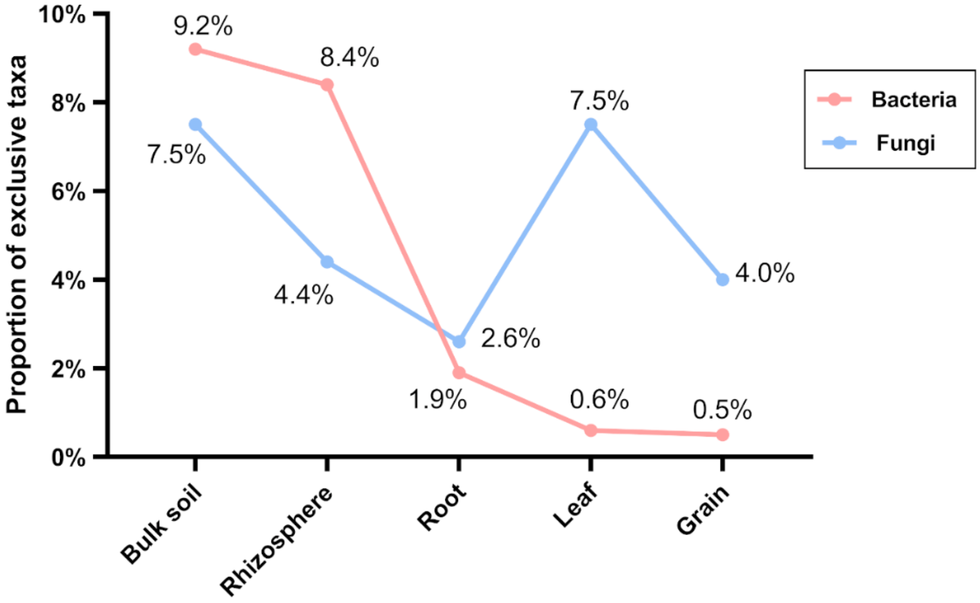


**Supplementary Figure S5:** Proportion of exclusive bacterial and fungal taxa per compartment.

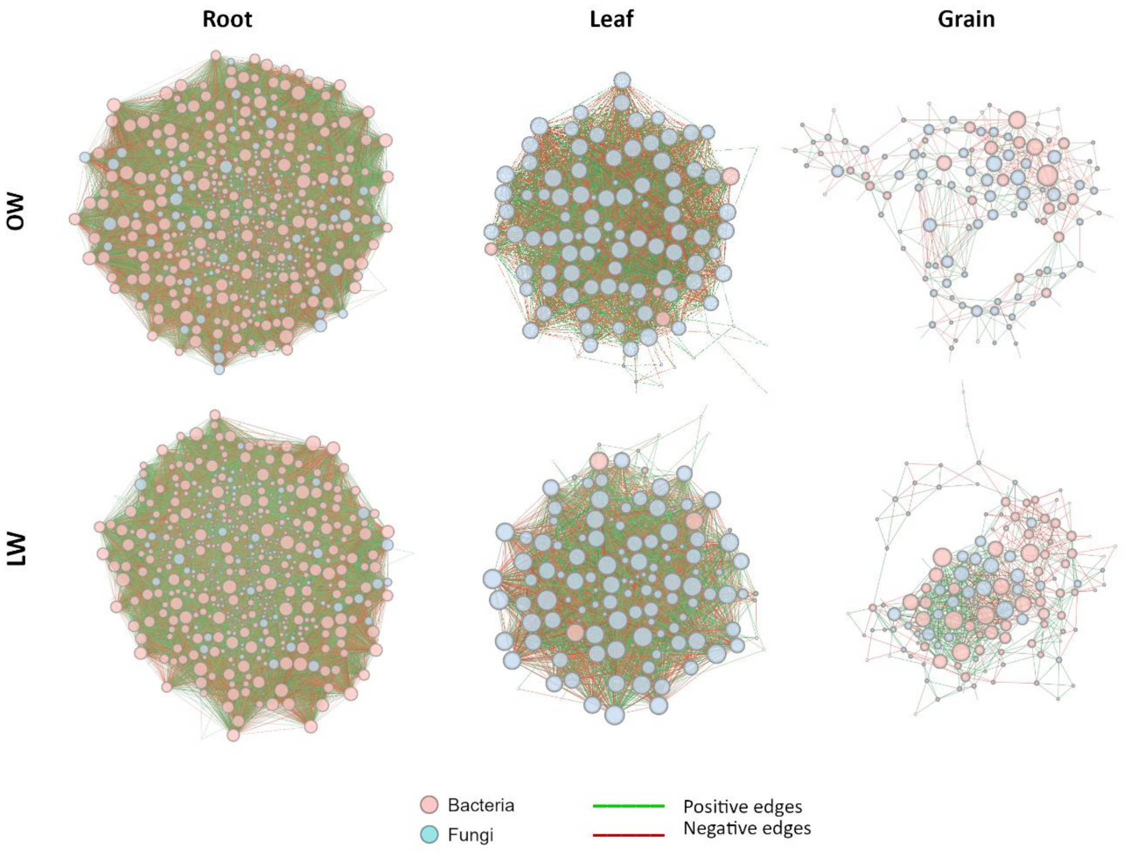


**Supplementary Figure S6: Microbial networks under different compartments and irrigation regimes.** Co-occurrence networks of plant compartments (root, leaf and grain) under optimal (OW) and limited (LW) watering. SparCC algorithm was used to calculate the network at ASV level with r > |0.6| and $P < 0.05$. Each node represents a single ASV. Pink nodes represent bacterial ASVs and blue nodes fungal ASVs. Node size is proportional to node degree within each network. Color of edges indicates the type of the interaction. Red, negative; green, positive.

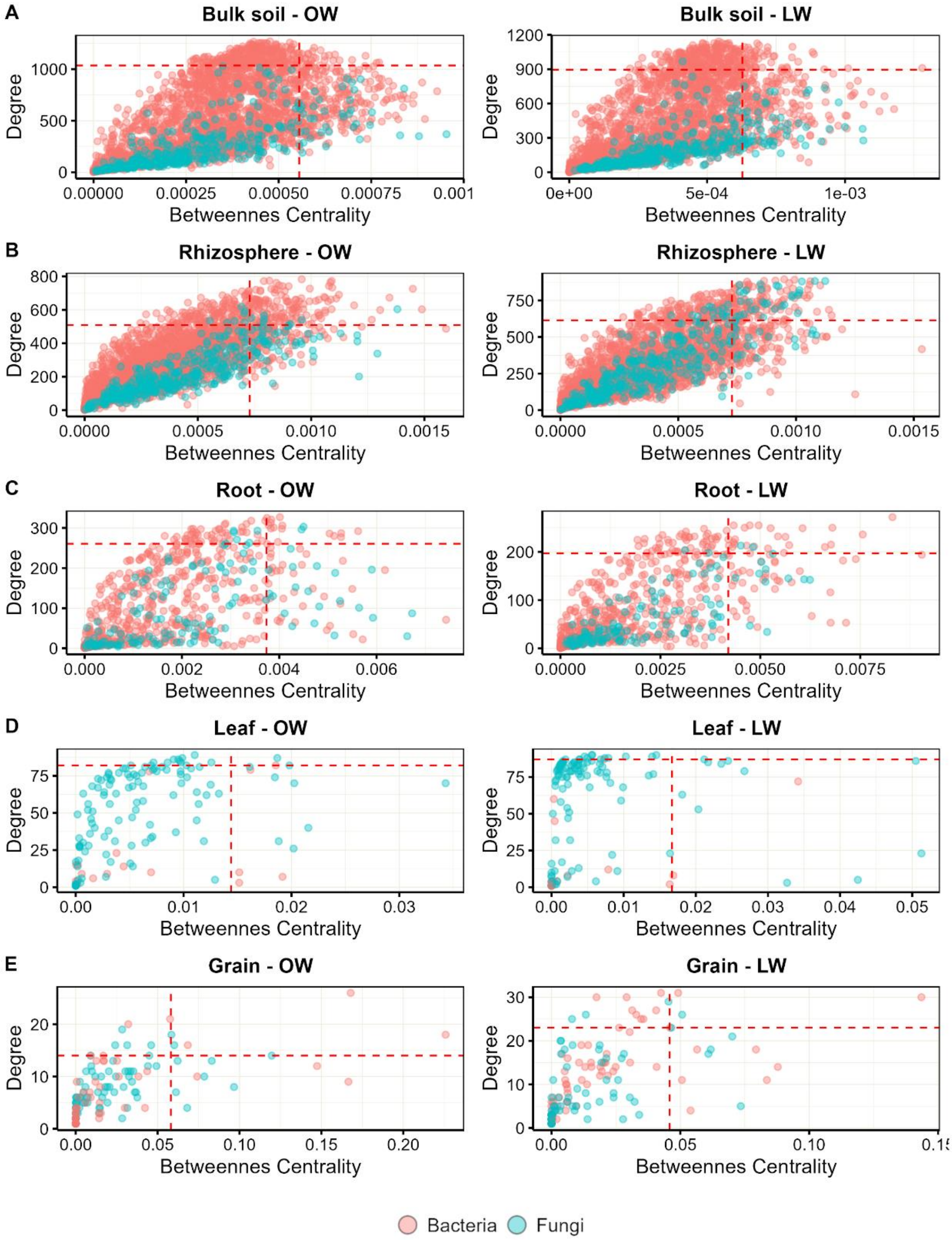


**Supplementary Figure S7:** Degree and betweenness centrality of bacterial and fungal nodes of (A) bulk soil, (B) rhizosphere, (C) root, (D) leaf and (E) grain compartments under optimal (OW) and limited (LW) watering. Red dash lines delimitate percentile 90th ($P_{90}$) in both axes.
